# Environmental DNA of aquatic macrophytes: the potential for reconstructing past and present vegetation and environments

**DOI:** 10.1101/2023.03.27.533457

**Authors:** Aloïs Revéret, Dilli P. Rijal, Peter D. Heintzman, Antony G. Brown, Kathleen R. Stoof-Leichsenring, Inger G. Alsos

## Abstract

Environmental DNA is increasingly being used to reconstruct past and present biodiversity including from freshwater ecosystems. Here, we first review and compare studies that use metagenomics, targeted capture, and various barcoding and metabarcoding markers, in order to explore how each of these methods can be used to capture aquatic vegetation diversity and change. We then investigate the extent to which such a record can be leveraged for reconstructing local environmental conditions, using a case study based on macrophyte ecological niches. We find that, with state-of-the-art DNA barcode reference libraries, using metabarcoding to target the P6 loop region of the chloroplast *trn*L (UAA) intron is optimal to maximise taxonomic resolution and the diversity of past macrophyte communities. Shotgun sequencing also retrieves a high proportion of aquatic macrophyte diversity, but has the lowest taxonomic resolution, and targeted capture needs to be more widely applied before comparisons can be made. From our case study, we infer past aquatic habitats from sedimentary ancient DNA records of macrophyte taxa. We reconstructed Holocene thermal range, continentality, water pH, trophic status, and light conditions in northern Fennoscandia. We show an overall stability since 9,000 years ago, even though individual lakes display different trends and variation in local climatic and physico-chemical conditions. Combined with the availability of near-exhaustive barcode and traits databases, metabarcoding data can support wider ecological reconstructions that are not limited to aquatic plant taxonomic inventories but can also be used to infer past changes in water conditions and their environmental drivers. Sedimentary DNA is also a powerful tool to measure present diversity, as well as to reconstruct past lacustrine and fluvial communities of aquatic macrophytes.

## 1 Introduction

Aquatic macrophytes are vascular plants that grow in lakes, ponds, streams, and wetlands. They are useful ecological indicators as their distributions are mainly determined by environmental factors at a local scale (Alahuhta et al., 2020; Johnson & Toprak, 2021; Poikane et al., 2018). Aquatic macrophytes play a major role in biogeochemical cycles in aquatic ecosystems and are therefore especially relevant for assessing the eutrophication of waterbodies (Penning et al., 2008) and waterways (O’Hare et al., 2018). Indeed, macrophyte communities effect the cycling of carbon (Reitsema et al., 2018), nitrogen (Dan et al., 2021; Zhang et al., 2021), and phosphorus (L. Wang et al., 2022) in both water bodies and their sediments.

Environmental DNA (eDNA) is a rapidly growing approach for addressing ecological and genetic questions about present and past organisms, from communities to populations. There is a need for more comprehensive monitoring of freshwater biodiversity (Harper et al., 2019), and a better understanding of the ecological responses of aquatic communities to climate shifts (Maasri et al., 2022). Compared to traditional methods of aquatic community monitoring and reconstruction (e.g. field survey, macrofossils, pollen), the eDNA techniques require less taxonomic expertise and sampling effort (Ji et al., 2021), and can provide higher temporal resolution (Buxton et al., 2018). For aquatic vegetation, eDNA may be comparable to, or even superior to, in-lake vegetation surveys (Alsos et al. 2018).

As a consequence of their biotope, aquatic macrophytes are readily incorporated into sediment, and are therefore especially well represented in sedimentary ancient DNA (*sed*aDNA) records (Alsos et al., 2022; Capo et al., 2021). Thus, combining a molecular palaeorecord of macrophytes with their known functional traits could allow for the reconstruction of past environmental conditions (Alsos et al., 2022; Dalla Vecchia et al., 2020; Dar et al., 2014). This has been implemented for a variety of animal and plant taxa using traditional methods, especially using macrofossil records. For instance, maximum summer temperatures in Fennoscandia throughout the Holocene have been reconstructed from chironomids (Brooks & Birks, 2000) or aquatic plant macrofossils (Väliranta et al., 2015). The latter can also help to reconstruct water level fluctuations (Väliranta et al., 2005). *Sed*aDNA has been shown to enhance such reconstructions, as it can provide higher consistency in detection of species than plant macrofossils (Alsos et al., 2016) and higher taxonomic resolution than pollen (Clarke et al., 2020; Courtin et al., 2021; Niemeyer et al., 2017), and can also be used as the sole proxy for the reconstruction of past terrestrial environments (Alsos et al., 2022). Therefore, it holds great potential for studies of past and present aquatic ecosystems (Stoof-Leichsenring et al., 2022), which would in turn expand the applicability of *sed*aDNA for abiotic reconstructions.

In addition to single-species barcoding assays, three major molecular approaches can be used to generate eDNA data, but they differ in taxonomic resolution and recovered richness estimates (Murchie et al., 2020). Shotgun sequencing is an untargeted technique (metagenomics), while (meta)barcoding and targeted capture enrich a part or parts of the genome of the taxa of interest (Box 1). Targeted capture is also called “hybridisation capture”, “capture probes”, or “target enrichment” in the literature. Markers used in plant barcoding and metabarcoding can be standard barcode regions like *rbc*L (654 bp) and *mat*K (862–910 bp); shorter ones developed to study degraded eDNA such as the P6 loop (51–135 bp) of the *trn*L UAA intron; or taxonomically broader loci not restricted to plants such as ITS and 18S (Hollingsworth et al., 2011). The presence of vascular plants detected in *sed*aDNA is often supported by other proxies such as present-day vegetation (Alsos et al., 2018), historical vegetation maps (Sjögren et al., 2017), macrofossils (Alsos et al., 2016), or pollen (e.g. Bjune et al., 2021; Liu et al., 2021; Parducci et al., 2019), thus confirming its reliability. However, an accurate detection of communities by eDNA heavily depends on the quality of the reference library (Y. Wang et al., 2021; Weigand et al., 2019), which should ideally be taxonomically exhaustive within the given geographic area. Other groups are likely to yield valuable information where the aquatic macrophyte signal is unavailable, thus increasing the value of palaeo-approaches for understanding both the past and the present (Brown, 2002). For example, Tyler and Olsson (2016) have also compiled the pH niches of Swedish bryophytes, which are a bycatch of the P6 loop. Using this marker in combination with both vascular plant and bryophyte traits databases could therefore give a very accurate depiction of pH evolution in the palaeo-record.

**Box 1:**
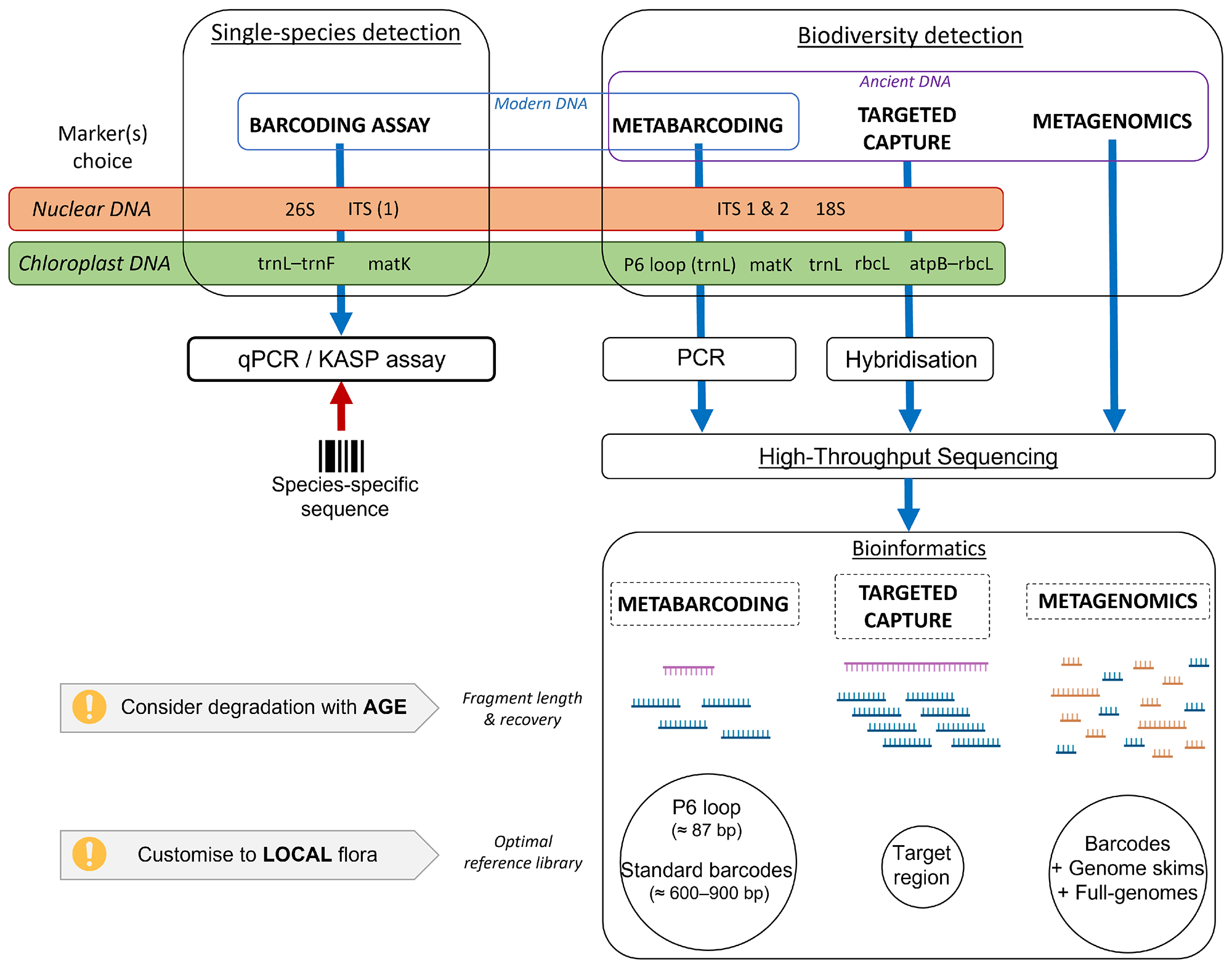
Methodological workflow of eDNA sample analysis for aquatic plants.

Here we review studies that identified aquatic macrophytes in ancient and contemporary lake sediments, soil, and water. To assess the potential of reconstructing past vegetation changes based on aquatic macrophyte community eDNA signals, we evaluate the taxonomic resolution and richness reported between the three major molecular approaches with a focus on ancient samples. We also show the potential for such reconstructions with an aquatic macrophyte case study, where we combined a *sed*aDNA metabarcoding dataset from ten lakes (Alsos et al., 2022) with ecological traits values from Tyler et al. (2021).

## 2 Materials and methodology

### 2.1 Compilation of aquatic macrophyte eDNA studies

We have compiled literature, including supplementary materials, which explicitly mentioned the detection of aquatic macrophytes using any eDNA method. We performed a literature search using the following phrases: *“sedimentary DNA aquatic vascular”*, *“sedimentary DNA macrophytes”*, *“freshwater environmental DNA plant diversity”*, *“sedimentary ancient DNA vascular”*, *“metabarcoding aquatic plants sediment”*, *“ancient sedimentary DNA vegetation”*, *“environmental DNA aquatic plant”*, *“environmental DNA macrophyte”*, and *“ancient DNA aquatic plant”*. For each search phrase, we scanned through the first 500 results for articles with pertinent features in the title or abstract (e.g. relevant method, field experiment, possibility of aquatic species, etc.), until we could compile 10 papers which met our criteria. We replicated this search on three search engines: first Google Scholar, then the bioRxiv preprint repository, and lastly the Web of Science Core Collection; we disregarded in our counts articles already recovered by previous search runs. In total, more than twelve thousand papers were scanned. We supplemented our search results with relevant studies from *AncientMetagenomeDir* (Fellows Yates et al., 2021), which collates metagenomics datasets, plus three studies mentioned in the bibliography of our review (Adame & Reef, 2020; Cannon et al., 2016; Schabacker et al., 2020), and relevant studies from the last *sed*aDNA review (H. Li et al., 2023).

When several studies investigated the same sites or an overlapping subset, we only considered the most recent study, unless an earlier study provided more data for aquatic macrophytes. We restricted our compilation to experiments carried out on field-derived samples that were recovered in natural conditions. We excluded marine studies of seaweeds and mangrove plants (Foster et al., 2021; Ortega et al., 2020), and *in vitro* and *in silico* studies (Du et al., 2011; Scriver et al., 2015). We also excluded non-vascular plant taxa, even when regarded as macrophytes by the authors (e.g. Characeae in Wang et al., 2021). We decided to disregard three diet studies where macrophytes were detected, as our focus is on water, soil, and sediment samples (canid faeces in Wood et al., 2016; human teeth in Ottoni et al., 2021, and in Sawafuji et al., 2020).

Using the resulting literature dataset, we compiled, where available: the number, type, and oldest age of eDNA samples, number of sites where aquatic macrophytes were detected through eDNA, methodological approach and loci targeted, DNA extraction method, number of PCR replicates, total and filtered count of DNA reads, counts of both raw and assumed true positive operational taxonomic units (OTUs) of all vascular plants, percentages of the latter assigned to the lowest taxonomic level (family/genus/species), number of OTUs assigned to aquatic macrophyte taxa, total number of aquatic macrophyte taxa retrieved from eDNA, counts from the latter assigned to the lowest taxonomic level, number, names, and versions of the reference libraries used for taxonomic assignment, identity cut-off for such assignment, and whether the study mentioned other proxies (i.e. pollen, macrofossils, and visual survey) (Supplementary Table S1). As OTU terminology contains synonyms, we collated figures reported as “sequences”, “OTUs”, “Molecular OTUs”, “Amplicon Sequence Variants”, and “taxa”. Note that several sequences may match the same taxon and therefore the latter measure is the most conservative in terms of diversity. We also compiled the geographical location of all sites with detected aquatic macrophytes through eDNA (Supplementary Table S2).

In order to consistently classify all reported vascular plant taxa as “aquatic” or not (Supplementary Table S3), we followed the functional groups given by the authors whenever possible. When references were conflicting (usually between one study identifying at the genus level and another identifying at the species level), we followed the original classifications for the respective studies, but these taxa were then ignored in other studies. All taxa found in the Swedish flora were classified according to our cut-off from Tyler et al. (2021), as described in the following section, and this classification always took precedence over categories given in the literature. The remaining taxa encountered were classified using the following floras: Flora of Colorado (Ackerfield, 2015); Calflora (Powell et al., 2022); Flora of Australia (ABRS, 2015); Flora of Tropical East Africa (Clayton et al., 1974); Flora Novae Angliae (Haines et al., 2011); Info Flora (Info Flora, 2022); Field Manual of Michigan Flora (Voss & Reznicek, 2012); Online Virtual Flora of Wisconsin (Wisconsin State Herbarium, 2022); PlantZAfrica (SANBI, 2022); eFlore (Tela Botanica, 2022); Svalbardflora (Elven et al., 2020); Flora of Victoria (Walsh & Entwisle, 1994); World Flora Online (WFO, 2022); and the following online databases: Global Biodiversity Information Facility (GBIF.org, 2022); Harrington et al., 2012; The Tree of Life Web Project (Maddison & Schulz, 2007); USDA Plants Database (USDA, 2022), all accessed in May 2022. Note that some ecologically broad genera, which include both terrestrial and aquatic species, such as *Carex*, *Equisetum*, *Juncus*, or *Ranunculus*, were included as aquatics only if originally defined as such by the study authors.

We assessed taxonomic resolution by method and marker used, in all vascular plants and in aquatic macrophytes, by computing the percentage of taxa identified to the lowest taxonomic level among family, genus, and species. Because these are compositional data, we applied a centred log ratio transformation ahead of further analyses. Barcoding assays were disregarded because they are species-specific: their taxonomic resolution would by definition be 100% at species level, with a single taxon detected (sometimes two were targeted). We also explored how total macrophyte richness correlates with macrophyte taxonomic resolution at each computed level and how richness and species-level resolution correlate with study year, number of samples, total sites, and age of the oldest sample; using generalised linear models (GLM, excluding outliers, Supplementary Figure S1). We assumed the distribution of errors to be respectively “Gaussian” and “Poisson” for continuous and discrete responses.

### 2.2 Northern Fennoscandia traits case study

Our case study focuses on northern Fennoscandia, as this region has the largest standardised lake *sed*aDNA dataset published to date, comprising eight PCR replicates each of 355 samples (Alsos et al., 2022; Rijal et al., 2021). In addition, an exhaustive DNA reference library (PhyloNorway) covers Norwegian vascular plants (Alsos et al., 2020a; Alsos et al., 2022), which maximises the chances of correct species identification and enhances the accuracy of subsequent analyses. A 100% match criterion was used for PhyloNorway, along with three additional reference libraries (ArcBorBryo, PhyloAlps, and EMBL-143; Alsos et al., 2022). A traits-value database for the Swedish vascular flora (Tyler et al., 2021) lists 30 parameters that can be used as ecological indicators. These traits represent both the abiotic and biotic components of niches (Ellenberg et al., 2001). We hereafter regarded “aquatics” as species that have a moisture requirement above level 9 (wet – temporarily inundated, e.g. *Caltha palustris*); on a scale from 1 (very dry) to 12 (deep permanent water). Thus, all our selected species are temporarily inundated (10, e.g. *Hippuris* spp.); live in shallow (< 0.5 m) permanent water (11, e.g. *Myriophyllum alterniflorum*), or deep permanent water (12, e.g. *Potamogeton praelongus*). As our taxonomic listing follows *sed*aDNA methodological constraints (i.e. some species are aggregated as they remain molecularly unresolved), taxa were attributed an averaged moisture value (e.g. 9.5 when one species is 9 and the other is 10; Alsos et al., 2022). Note that this scale emphasises water depth rather than plant growth form (free-floating, submerged, floating-leaved, emergent, or marginal), as used in some studies (e.g. a variant in Tyrrell et al., 2022) (Supplementary Table S4). Depths of the lakes studied ranged from 1.2 to 34.8 m, most of them being between 4 and 15 m (see supplementary material in Rijal et al., 2021).

A *sed*aDNA plant metabarcoding dataset spanning across 10 lakes in northern Fennoscandia (Rijal et al., 2021) with taxonomic assignments updated by Alsos et al. (2022), was further subsetted to retain only aquatic macrophyte taxa, by comparing the recorded species list with their respective moisture requirement (Tyler et al., 2021), following the standardisation of taxonomic nomenclature in Alsos et al. (2022). The majority of the data were initially published in Rijal et al. (2021), with a focus on how climate and soil nutrients affected overall taxonomic richness over the last twelve millennia. Alsos et al. (2022) expanded the dataset and revised the sequence identification using the new PhyloNorway DNA reference library to reconstruct post-glacial establishment across sixteen millennia, and then investigated traits related to colonisation, such as dispersal mode and pollinator dependence. The scope of the present case study is to carry out a detailed examination of aquatic plant taxa, to support the proper reconstruction of major abiotic variables important for aquatic macrophytes distribution.

The final reconstructed six traits were heat- and cold requirements, continentality, pH, nitrogen availability, and light optimum. Most of these factors are known to be drivers of aquatic macrophyte distribution patterns (Dar et al., 2014). Out of the 30 total traits from Tyler et al. (2021), we firstly discarded ten traits with less than a minimum of seven distinct trait values across all aquatic taxa; thus eliminating nitrogen fixation (1 level), assumed immigration time to Sweden (1), parasitism (1), carnivory (2), mycorrhiza (2), seed dispersal (3), pollinator dependence (4), seed longevity (5), nectar production (5), and tolerance to grazing (6). We additionally discarded three traits that had missing values for seven or more aquatic taxa (soil disturbance, phenology, seed dormancy); and six that were irrelevant or marginal for our environmental reconstruction (occurrence in Sweden, red-listing, photosynthetic pathway, invasiveness, longevity, vegetation type). Thus, the 11 remaining ecological traits to be tested for environmental reconstruction were: biodiversity relevance (log of trophic associations), cold- and heat requirements, temperature optimum (computed from cold and heat requirements), continentality, light optimum, moisture, pH, nitrogen and phosphorus availabilities, and salinity. We performed a Spearman’s rank correlation analysis of these variables to exclude collinear explanatory variables: one of two traits was discarded whenever their paired correlation coefficient value (r) was > 0.7 (Supplementary Figure S2). Such high correlations were found between cold requirement and temperature optimum (r = 0.937), pH and salinity (r = 0.770), and nitrogen and phosphorus requirements (r = 0.746); and we kept the formers. Moisture and biodiversity relevance were dropped to simplify graphical representation, as they were less informative than other retained traits.

We investigated the possibility of reconstructing past environmental changes by looking for major shifts among traits values; both at the regional scale, combining data from all 10 lakes, and the local single-lake scale. For the regional dataset, we aggregated samples by 500-year time slices and plotted the distribution of trait values through time, using their respective proportions across all eight PCR replicates. For individual site reconstructions, we did not merge samples into time slices before plotting the distribution of traits values as their respective proportions across replicates through time.

Additionally, we checked for potential correlations between species traits and their first recorded and estimated arrival dates in the region (Alsos et al., 2022), to see if some traits could explain the arrival of aquatic macrophytes. We further evaluated how the composition of aquatic taxa was affected by our six selected traits, using multivariate analysis. First, we ran a detrended correspondence analysis (DCA) as a preliminary analysis to select an appropriate multivariate analysis. As the gradient length was short (0.76), we applied a linear ordination method, namely a redundancy analysis (RDA), with presence/absence data as the response variable. All analyses were run in R v4.1.1 (R Core Team, 2022).

## 3 Results

### 3.1 Published data compilation

We found 62 studies that explicitly mentioned the detection of aquatic macrophytes using eDNA, encompassing 450 sampling sites (Supplementary Tables S1 & S2). The majority of sampling sites originate from boreal, arctic, or high-altitude bioclimatic zones, especially for studies of ancient DNA (Figure 1). Our review covers 4,126 samples in total, 1,124 of which are modern samples only (water, lake or river surface sediment, or undated but assumed to be recent material). The majority of studies (55%) target lacustrine or fluvial sediment, including thermokarst lakes (thus the permafrost active layer). Several studies reviewed here also used water samples (26%) from lakes and rivers, soil samples (19%) encompassing permafrost sedimentary complexes and subfossil stream deposits from river bluffs, and peat samples (8%). Four studies used samples of different types. Lastly, despite our selection criterion, two studies also included marine sediment (Figure 1): these sites are a tidal basin which later turned into a freshwater lake due to isostatic rebound (lake Nordvivatnet, see Brown et al., 2022), and a pre-transgression terrestrial area near the past coastline (Gaffney et al., 2020).

**Figure 1:**
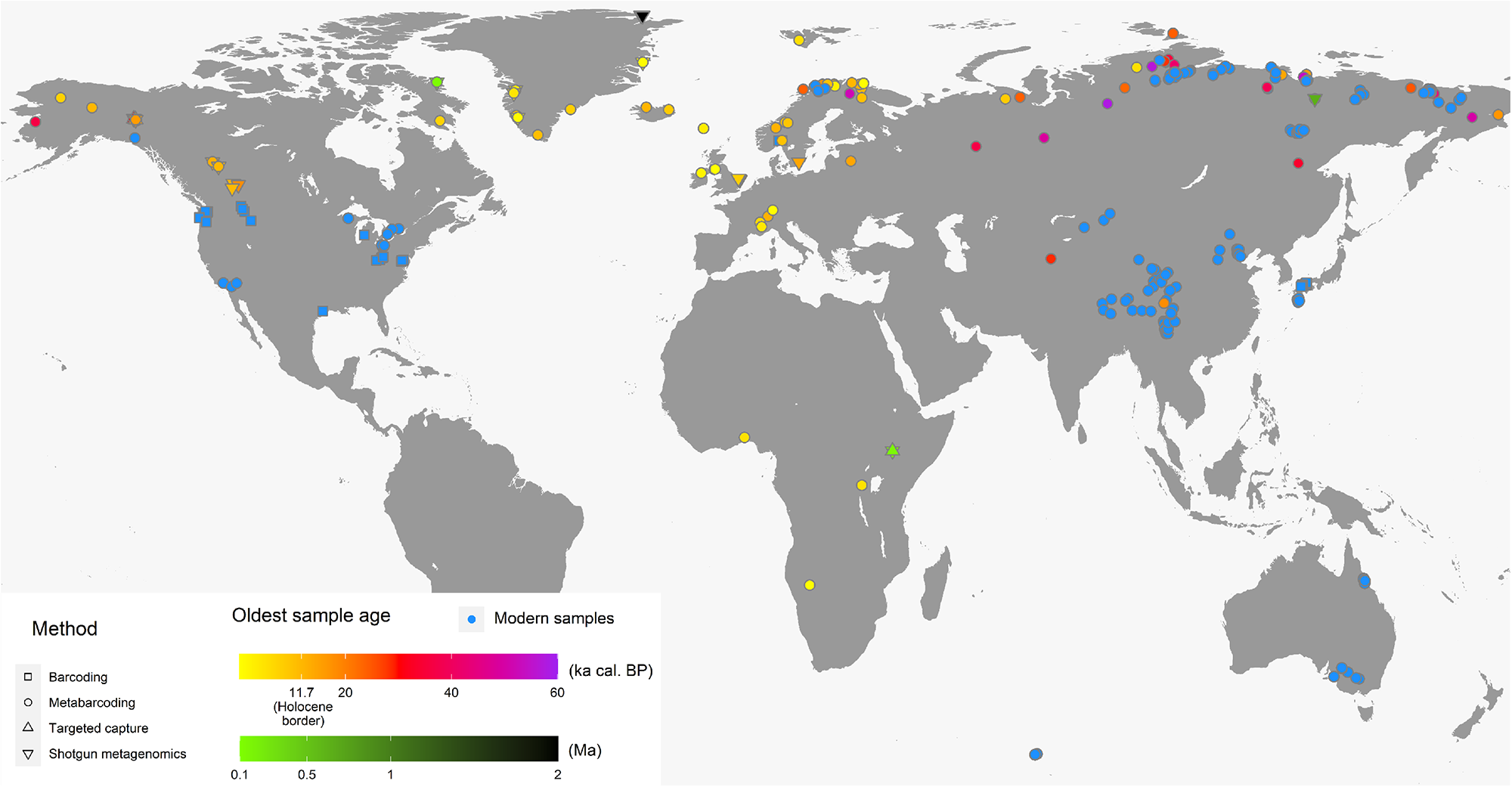
World map showing the location and age for eDNA studies reporting aquatic macrophytes. Sites in blue account for modern samples, i.e. water, surface sediment, and other undated material. All studies sampled freshwater or terrestrial material, except for two which retrieved marine sediment.

Our review is dominated by ancient DNA (aDNA) studies (*n* = 41) compared to modern eDNA studies (*n* = 21), although the former often include modern samples through analysis of surface layers in sediment core records. There are three main approaches to eDNA analyses (Box 1) and most of the 62 studies reviewed used only one of them, while seven combined several methods, at least on a subset of their samples. Metabarcoding is used in most studies (*n* = 49), followed by shotgun sequencing (*n* = 10), barcoding assays (*n* = 9), and targeted capture (*n* = 2). Among barcoding and metabarcoding studies, seven targeted multiple loci. Sample age is found to be strongly related to the choice of methods and markers: all shotgun sequencing studies were performed on ancient samples, whereas all barcoding assays focused on modern material (Table 1).

**Table 1:**
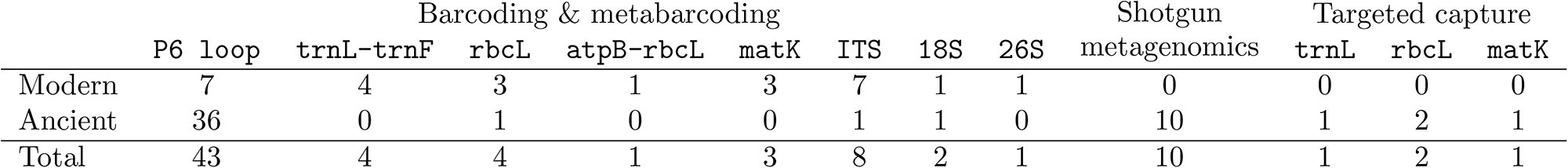
Number of reviewed publications by method used and locus targeted, depending on their sample age. Studies using several methods or targeting several markers are counted multiple times.

In total, 319 aquatic plant taxa were reported (Supplementary Table S3). However, 50 of these taxa were excluded because they i) are not vascular plants (*n* = 22), ii) were erroneously classified (*n* = 3), iii) had an invalid taxonomic name (*n* = 1), iv) were assumed false positives as they were outside of their continental biogeographical range (*n* = 6), or v) were only detected by methods other than eDNA (*n* = 18). Thus, 269 aquatic taxa belonging to 45 families, 118 genera, and 196 species were used in this review (Supplementary Table S3). There was a clear difference in approach between ancient and modern samples (Tables 1 & 2). Shotgun sequencing has only been applied to ancient samples, whereas barcoding assays have only been used to detect single species in modern water samples, for the monitoring of invasive or threatened species. Only two studies used targeted capture (Table 2); and this was to investigate plant and animal diversity in permafrost between 10,000 and 30,000 years old (Murchie et al., 2020), and in Late Pleistocene sediment from an arid palaeolake basin (Krueger et al., 2021). Metabarcoding has been applied to both modern and ancient samples, using multiple markers. The *trnL* P6 loop is by far the most used marker for ancient samples, having been used in 92% of all aDNA metabarcoding studies reviewed (Tables 1 & 2). For modern samples, a much wider array of markers is used including the standard barcodes *rbcL* and *matK* (Table 1).

**Table 2:**
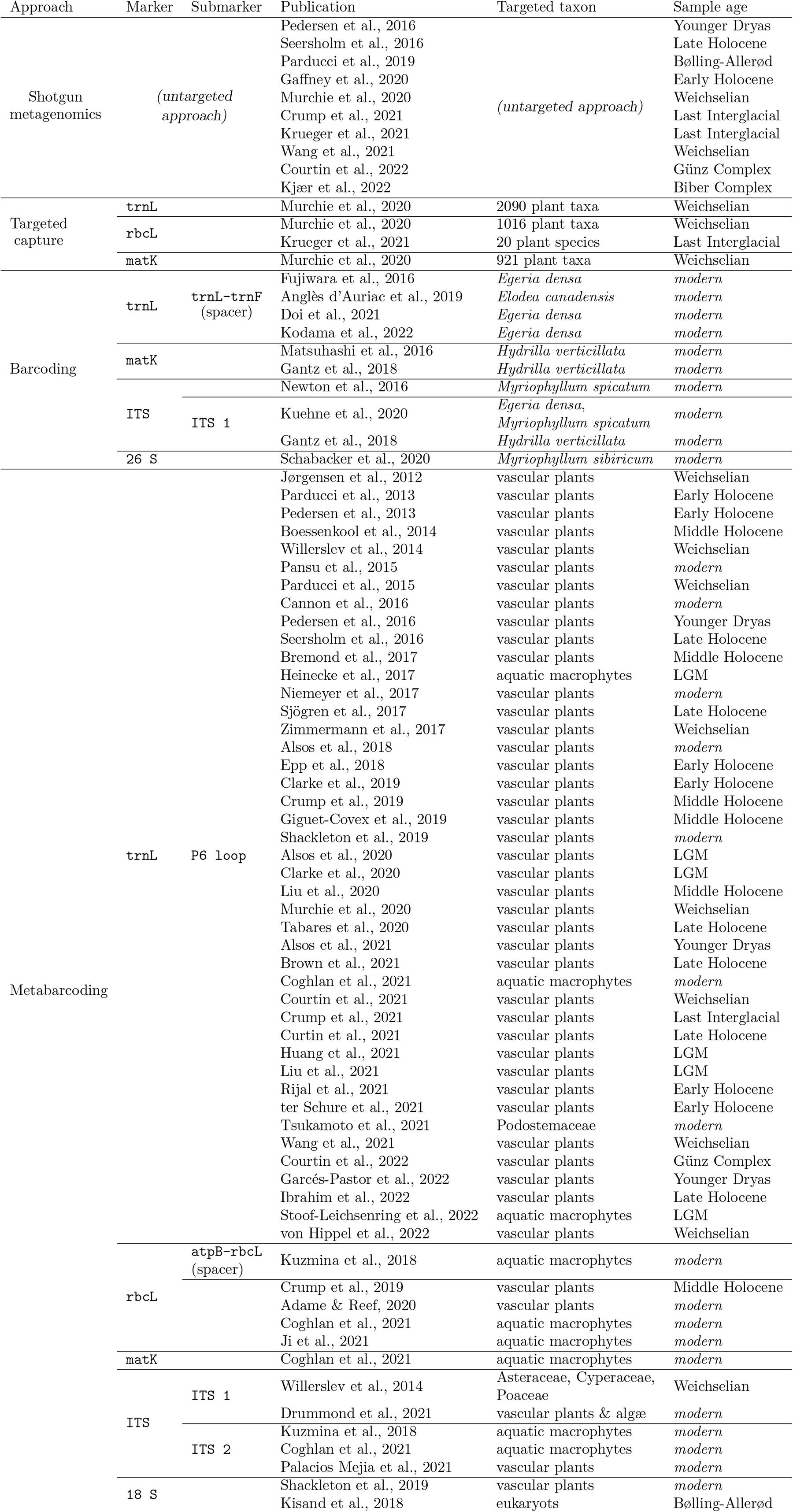
Summary of publications which reported detection of aquatic macrophytes in eDNA, their molecular methodologies, aimed taxonomic breadth (within Viridiplantae), and stadial/interstadial corresponding to the oldest sample. LGM is Last Glacial Maximum, Bølling-Allerød (14.7 to 12.9 ka), Younger Dryas (12.9 to 11.7 ka), Early Holocene (11.7 to 8.3 ka), Middle Holocene (8.3 to 4.25 ka), and Late Holocene (4.25 to 0.0 ka).

The earliest study to report aquatic macrophytes detected in eDNA was published in 2012 by Jørgensen et al., which found *Caltha* sp. and *Pleuropogon sabinei* in Weichselian sediments from Lake Taimyr, Siberia. Indeed, the occurrences of macrophyte eDNA were exclusively from metabarcoding of ancient material, until 2016 (Figure 2). The first studies to use multiple eDNA detection methods at once were published in 2016, introducing metagenomics as a new tool to study macrophyte palaeodiversity and inaugurating a more active period in this field (Figure 2). Between 2016 and 2022, barcoding assays have been implemented to monitor the invasive species *Egeria densa*, *Hydrilla verticillata*, *Myriophyllum spicatum*, *Elodea canadensis*, and *Myriophyllum sibiricum*. A second increase in publications is observed from 2020 onwards, with (to date) the recent detection of ancient macrophyte DNA in samples over two million years old (Figure 2).

**Figure 2:**
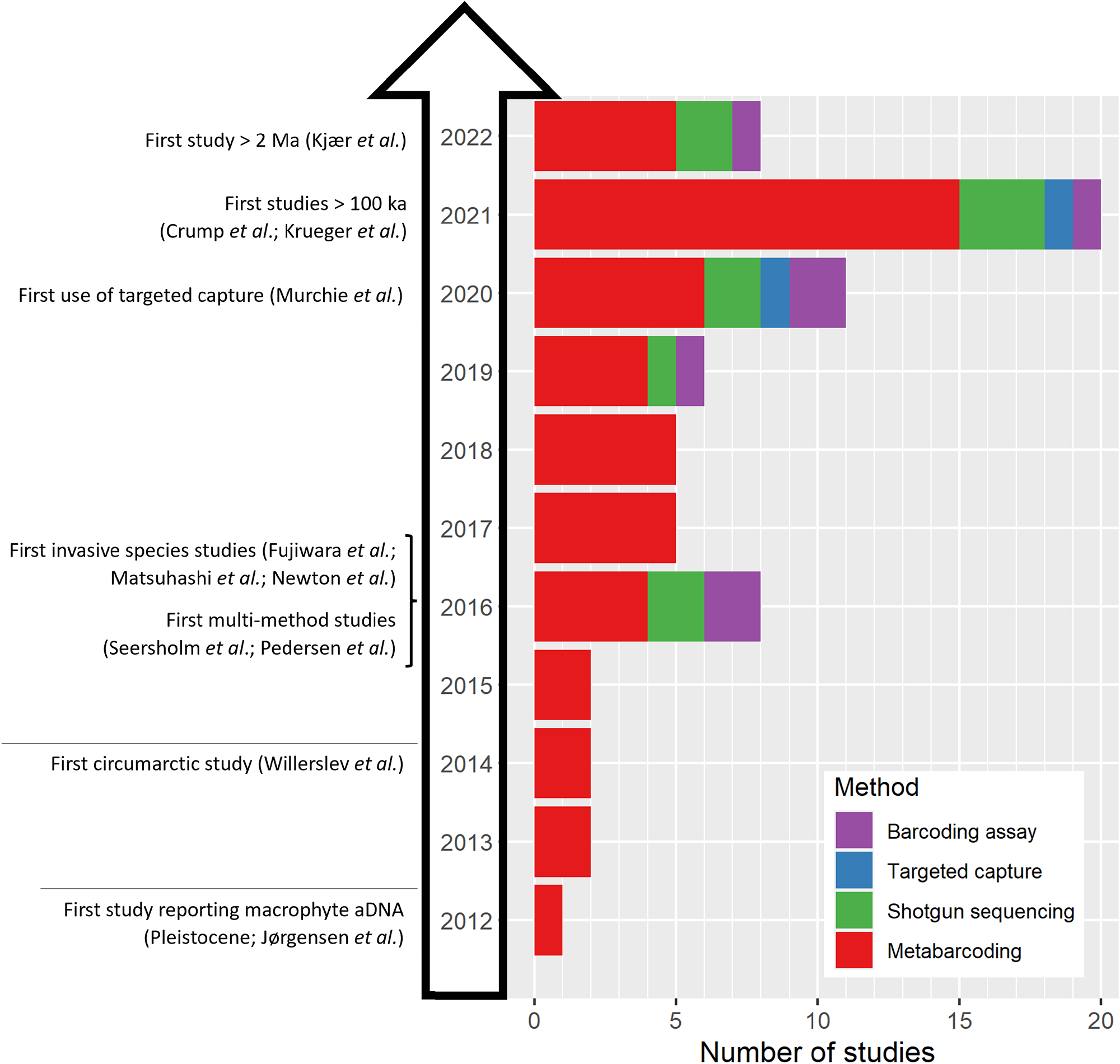
Number of reviewed studies per publication date and approach used. Publications featuring several methods are shown as multiple studies.

Sedimentary DNA has been successfully used to unravel past changes in macrophyte communities at various timescales. For instance, Bremond et al. (2017) who reconstructed plant palaeocommunities in a tropical lake over the last 5,000 years found aquatic taxa which ecologically corresponded to the seasonal flooding of the shores. They also detected the common water-hyacinth *Pontederia crassipes*, a recently introduced plant now dominating the vegetation of the shallows. On a longer time scale, Alsos et al. (2022) reconstructed the postglacial arrival of aquatic macrophytes in northern Fennoscandia, and although it was the last growth form to appear last (ca. 12,900 cal BP) nearly all taxa appeared at once, highlighting their capacity to immigrate rapidly once suitable environmental conditions are established. Ibrahim et al. (2022) detected the presence of the alien Canadian waterweed *Elodea canadensis* in *sed*aDNA over the last centuries, and Anglès d’Auriac et al. (2019) tracked it from water samples over a narrow spatial scale. To date, no study has addressed evolutionary questions for macrophytes with *sed*aDNA, but examples on other plant taxa have opened a way.

Many studies referred to one, or more, other proxies in order to confirm their DNA findings, but only few carried out an actual comparison with their DNA samples while the rest referred to previous work from the site. Papers investigating modern communities often chose to carry out vegetation surveys. Such surveys often only target a specific taxon of interest in the vicinity (e.g. Tsukamoto et al., 2021), a list of species detected at the spring (Palacios Mejia et al., 2021), or in a given area (e.g. Drummond et al., 2021), or a more complete version providing abundance and biomass estimates (e.g. Ji et al., 2021). The latter showed that eDNA metabarcoding of a ca. 300 bp barcode was better than a visual survey for submerged vegetation, which is often hampered by factors such as waterbody area, flow speed, turbidity, and depth (J. Hughes et al., 2018). Moreover, Alsos et al. (2018) demonstrated that lake sediment eDNA from recent samples largely matches the extant vegetation with 88% of dominant and common taxa and 60% of rare taxa detected, and additionally records taxa missing in the survey that are likely true positives such as plants growing in deep water. Detection is also highly related to environmental conditions, especially temperature and water conductivity (Stoof-Leichsenring et al., 2022). Macrophyte communities constitute a good indicator of environmental conditions, especially in regions with high aquatic plant diversity which allows sensitivity to the spatial heterogeneity of the ecosystem, and can distinguish land use in river sections (Ji et al., 2021). Focusing on this group in *sed*aDNA could support a temporal reconstruction of water quality and trophic status, for instance where human-induced eutrophication is a major stake. If multiple sedimentation basins are encountered, a spatial approach could even be added, as showcased by Ibrahim et al. (2022) with the overall vegetation in Lake Constance. Pollen remains a widely used proxy for comparison to plant *sed*aDNA, but no comparison has focused on aquatic taxa which, in general, have more locally dispersed pollen (by water or insects) but poorer taxonomic resolution than the P6 loop (Sønstebø et al., 2010). Macrofossils only identify a small fraction of the plant community but are particularly suited for aquatics (Parducci et al., 2019).

We found taxonomic resolution to vary greatly between methodological approaches (Figure 3). Metabarcoding has contrasting results that are dependent on the selected marker. The P6 loop identified on average 55% of all aquatic taxa detected to species-level (Figure 3, Supplementary Table S1). ITS 1, ITS 2, and *rbc*L have higher discriminative power, and respectively assigned 74%, 79%, and 80% of aquatics to species; although each of them was used in only two to four studies (Supplementary Table S1). Shotgun sequencing identifies 12% and 45% of aquatic taxa respectively at family and genus level, but only 38% at the species level (Supplementary Table S1). For metabarcoding, the taxonomic resolution was generally higher for aquatics than for other vascular plants (Supplementary Table S5). Regarding targeted capture, Murchie et al. (2020) created a bait-set targeting the three chloroplast loci *trn*L, *rbc*L, and *mat*K based on sequences from 921–2090 arctic and subarctic taxa (Table 2), but the resulting sequences could not be identified beyond genus level (Figure 3). On the other hand, Krueger et al. (2021) targeted *rbcL* only with a bait-set designed from just 20 vascular plant species including 5 aquatics (Table 2), and reported 6 aquatic species and 1 genus (Figure 3, Supplementary Table S1).

**Figure 3:**
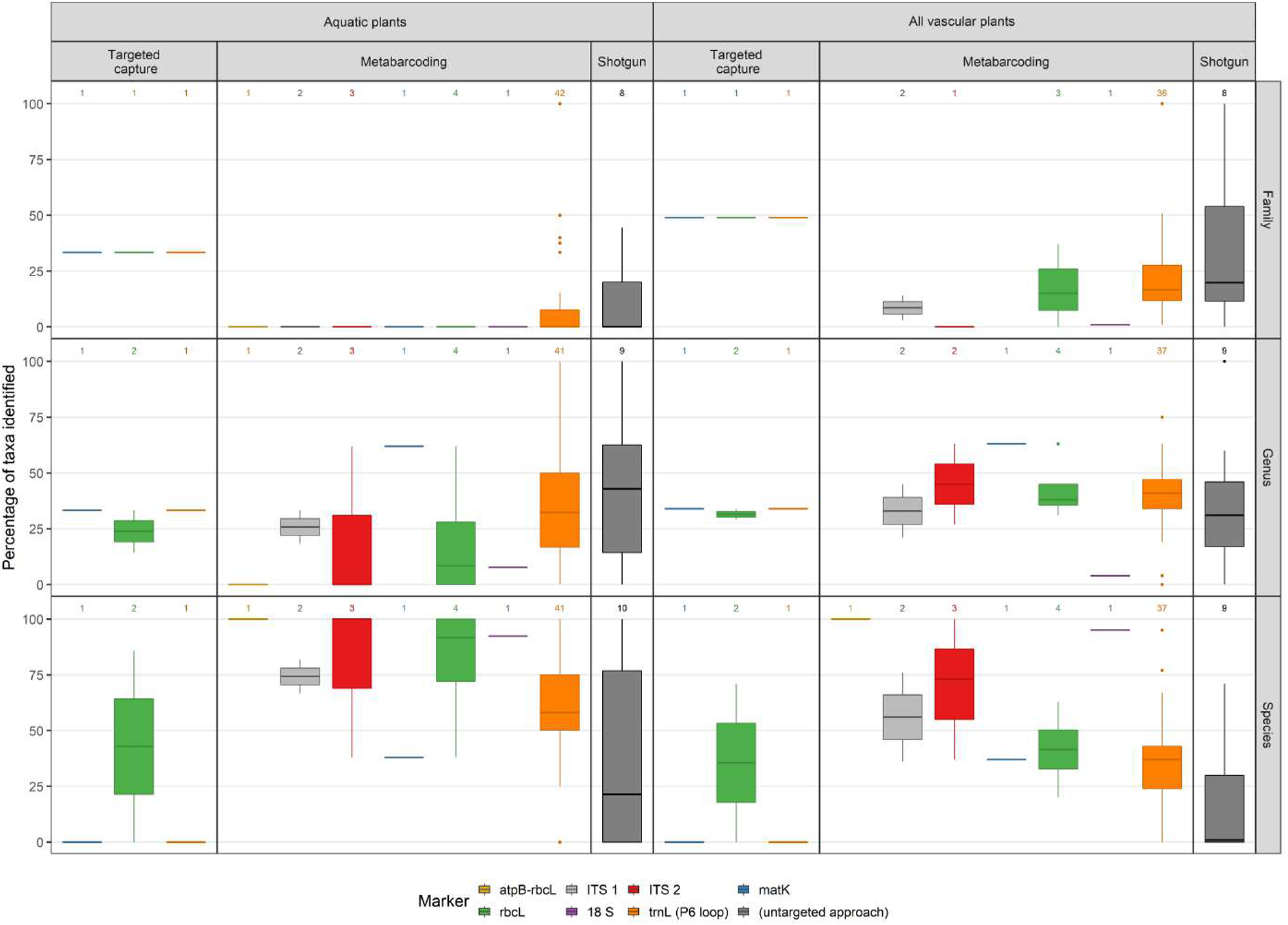
Taxonomic resolution in percent of taxa identified to the family, genus, or species level; by method and marker used, for aquatic macrophytes only (left) and all vascular plant taxa (right). Sums of taxa identified might not reach 100%, i.e. taxa were identified at a level higher than family. Number of studies is shown above each boxplot, it is inconsistent whenever a study gives partial results and mentions only a certain taxonomic level. We excluded datasets which used single-species approaches, because by definition their taxonomic resolution would be 100% at the species level.

The quality and exhaustiveness of the reference libraries affects the taxonomic resolution for all three methods (Box 1). The reviewed studies used a median of two reference libraries to assign their OTUs (Supplementary Table S1): they usually used a global one, such as GenBank or EMBL, and a better curated one with focused regional coverage, which reduces the possibility of sequence sharing with geographically non-plausible species (e.g. PhyloNorway, ArcBorBryo).

The aquatic macrophyte richness is not correlated to their taxonomic resolution at the species level, implying that a method with higher taxonomic resolution does not result in a higher diversity captured (Table 3). However, total richness detected depends in turn on other parameters: it is higher in more recent publications and may also decrease with (oldest) sample age especially for P6 loop studies (Table 3, Supplementary Figure S1), but old (> 60,000 cal BP) samples are too scarce to be considered representative. However, richness detected is also largely affected by DNA preservation, itself depending on sample age as well as environmental factors (Jia et al., 2021).

**Table 3:**
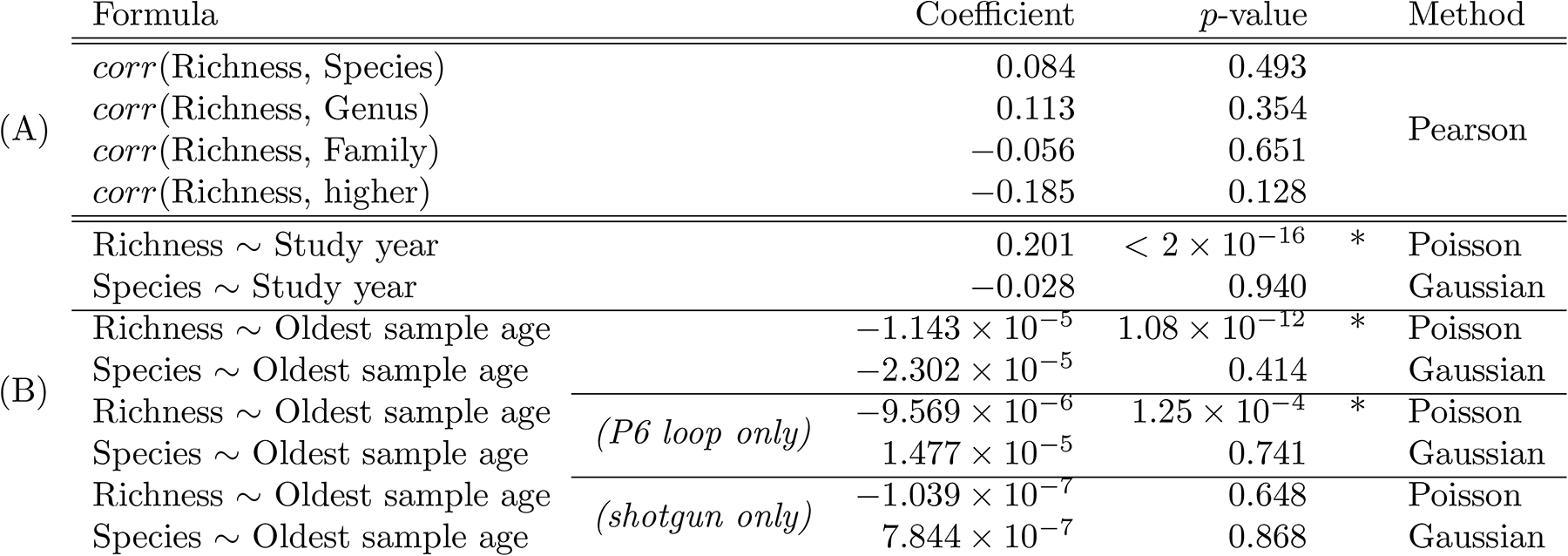
Parameters and output of (A) correlation tests between macrophyte diversity detected and resolution at four taxonomic levels after a centered log ratio transformation, and (B) Generalised Linear Models assessing the respective influences of the year of study and the age of samples on macrophyte diversity detected, and on its species-level resolution. We used a Poisson distribution in the model as the response variable was a count data. Asterisks indicate a significant (p < 0.05) relationship. Values were rounded to three decimal places, and outliers were removed.

### 3.2 Case study: reconstruction of past aquatic conditions in 10 lakes from northern Fennoscandia

There were 28 aquatic taxa in our northern Fennoscandia case study, 15 of which were unambiguously identified to the species level. In total 19 taxa had trait values available across all six reconstructed traits, i.e. heat- and cold requirements, continentality, pH, nitrogen availability, and light optimum. We found no significant correlation between the first recorded or estimated arrival dates of taxa and our selected traits, correlation coefficients being between – 0.3 and 0.5 for the detected arrivals, and between – 0.4 and 0.1 for the estimated arrivals. The first and second RDA axes respectively explained 17.7 and 6.4% of the variation in aquatic macrophyte community composition (Figure 4). Of the six traits included in the RDA, heat requirement had the highest correlation with axis 1 (ρ = – 0.974), and continentality had the highest correlation with axis 2 (ρ = 0.804).

**Figure 4:**
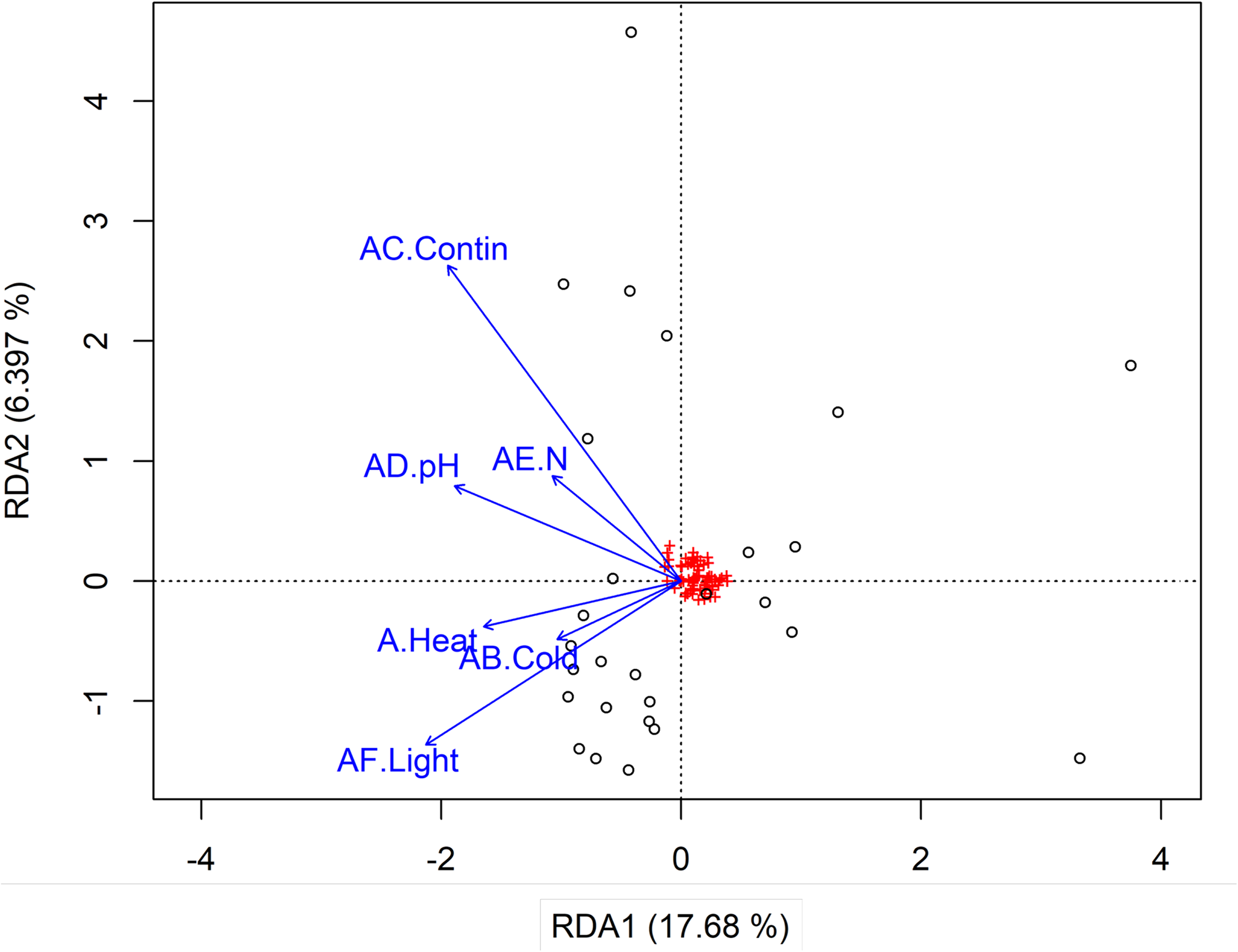
Redundancy analysis (RDA) of macrophyte community composition constrained by 6 selected environmental traits. Percentages of variance explained by each axis are given in brackets.

Our reconstruction of past environmental conditions across northern Fennoscandia shows the variability in the diversity of niches, and an overall stability since 9,000 calibrated years before the present (cal BP) (Figure 5). There is a weak, but steady acidification trend: the early record is largely dominated by circumneutral (pH ∼6.5) taxa, but modern communities are characterised by moderately acidic (pH ∼5) taxa. Since the end of the Early Holocene, heat requirement, continentality, nitrogen availability, and light optimum are dominated by a single trait value representing over half of weighted PCR replicates. Indeed, since ca. 8,000 cal BP, the majority of niches realised by aquatic macrophyte communities has been characterised by a low heat requirement (i.e. value 3 = reaching as high as the low-alpine belt, *sensu* Tyler et al. (2021)), with no effect of continentality on their geographical range (i.e. values 4.5–5 = distributed indifferently across Scandinavia, *sensu* Tyler et al. (2021)), moderately nitrogen-poor water, and half-shade conditions. However, the pre-Holocene record is not a fully regional signal, as the Fennoscandian ice sheet still covered most of the region at 12,000 cal BP (A. Hughes et al., 2016). Only lakes Sandfjorddalen and Nordvivatnet have aquatic macrophyte records predating the Holocene (> 11,000 cal BP). Although the Langfjordvannet core extends to 16,000 cal BP, no aquatics are detected before 10,000 cal BP. It is worth noting that this lake is the deepest (34.8 m) thus macrophytes cannot be found in its centre, as their maximum growth depth is about 12 m (Sheldon & Boylen, 1977). From 10,000 cal BP onwards, there are more than five lake records with aquatic macrophytes (Supplementary Figure S3). Species requiring the warmest conditions, such as *Cicuta virosa* and *Stuckenia vaginata* (Supplementary Figure S4), disappear between 5,000 and 3,000 cal BP (Figure 5), which can be interpreted as lower summer temperatures. This pattern is consistent with the long-term cooling trend after the mid-Holocene thermal maximum; but the late reappearance of *Stuckenia vaginata* in lakes Gauptjern and Sandfjorddalen at ca. 2,700 cal BP, then in lake Horntjernet at 100 cal BP indicates a recent warming (Supplementary Figures S6a, S6b, & S6c). Pioneer aquatic communities developed under intermediate light conditions, with most taxa requiring sun but enduring some shading, and there was a progressive shift towards increased heterogeneity between 9,000 and 4,000 cal BP. The modern presence of half-shade tolerant taxa as well as taxa requiring full sunlight exposure, reflects an increased habitat complexity and/or the development of closed-canopy riparian forest. The slow progress of light-dependent taxa could be linked to winter ice dynamics, and be indicative of winters becoming progressively shorter.

**Figure 5:**
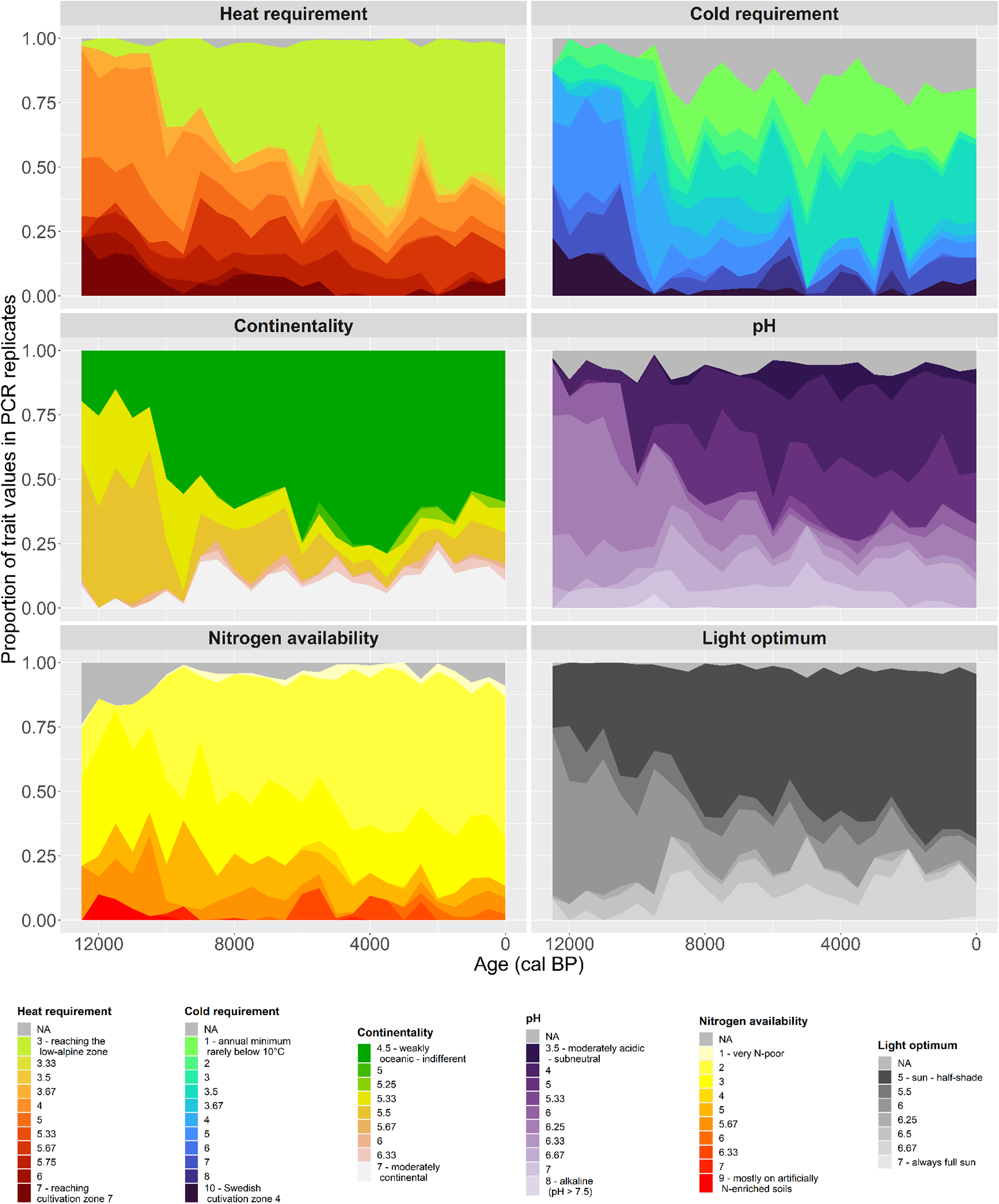
Proportion of traits values (see methods) in PCR replicates across northern Fennoscandia (all 10 lakes) through the end of Late Pleistocene (12,500 to 11,700 cal BP) and the Holocene (11,700 cal BP to present), in 500-years bins. A high trait value indicates a strong expression of the trait.

Individual lakes display different trends in environmental reconstructions, depending on their community composition and turnover. We retained only those lakes which have an average richness over time of five or more aquatic taxa (Sandfjorddalen = 8.6, Nordvivatnet = 6.2, Sierravannet = 5.6, Horntjernet = 5.6, Kuutsjärvi = 5), so that the distribution of traits values is robust enough to support environmental inferences for a single site. For instance, lake Horntjernet sees a slow decrease of cold-tolerant taxa, followed by an absence around 4,000 cal BP then an increase (Supplementary Figure S5a), with higher proportions of *Ranunculus* cf. *reptans* in PCR replicates (Supplementary Figure S6c). Lake Sandfjorddalen, which has a record that spans the entire Holocene (except for a hiatus between 5,500 and 3,000 cal BP, Supplementary Figure S6b), shows long-term stability in environmental conditions (Supplementary Figure S5b), partly due to a low compositional turnover (Supplementary Figure S6b). Such consistency suggests that the lake depth has probably remained stable and shallow (presently 1.23 m) throughout, because we would expect a complete drying or an increase in water level to cause significant changes in the macrophyte community. Conversely, lakes Horntjernet and Kuutsjärvi display more temporal variability in reconstructed environmental conditions (Supplementary Figures S5a & S5c). There is a steady decrease in continental taxa over the long term in Kuutsjärvi; whereas Hortjernet is dominated by moderately continental taxa until ca. 9,000 cal BP, and continentality increases again from 3,500 cal BP. Initial conditions in Horntjernet were circumneutral to slightly acidic until 8,500 cal BP, when pH drops below 6.5 concurrently with the arrival of *Pinus* forest (Rijal et al., 2021). The reappearance of *Stuckenia vaginata* in the most recent samples indicates another shift towards a more neutral pH, as this species has a pH tolerance of ∼7. *Cicuta virosa* appears in Kuutsjärvi as early as 9,000 cal BP and disappears four millennia later (Supplementary Figure S6d); its presence indicates eutrophication and warmer temperatures.

## 4. Discussion

The general hypothesis that the DNA methodology used will affect the reconstructed aquatic plant assemblage, including estimated diversity, composition, and richness, is supported and the degree of variation is showcased by this study. We further demonstrate that the reconstruction of past environmental conditions based on the aquatic macrophyte *sed*aDNA signal is possible at both local and regional scales.

### 4.1 Global distribution of reviewed studies

Most sites concentrate in high-latitude or high-altitude regions (Figure 1). Polar regions are especially vulnerable to invasive species and so will be a priority for eDNA-based aquatic biodiversity monitoring, and there are also comprehensive DNA reference libraries available for this region for the P6 loop (Alsos et al., 2022; Sønstebø et al., 2010; Willerslev et al., 2014), standard barcodes (Alsos et al. 2020; Kuzmina et al., 2017), and genome skims (Wang et al., 2021). However, genetic reference libraries are not evenly available across the globe, and their existence and implementation depend on the diversity and taxonomic specificities in each freshwater ecoregion. Aquatic macrophytes are less diverse and have larger ranges in northern Fennoscandia as compared to intertropical zones (Murphy et al., 2019), which contributes to the near-exhaustivity of the PhyloNorway reference library. In contrast, the large reference database used by Ji et al. (2021) to detect over a hundred macrophyte species along the Chaobai river in China, was based on the global Nucleotide Sequence Database from NCBI’s GenBank, which suffers from a lower curation rate and may not be fully representative of the local flora. Additionally, using such libraries with global coverage can match genuine local sequences to their exotic counterparts, but this source of error can be avoided with a stringent cross-checking of the resulting taxa list (Supplementary Table S3, see taxa excluded from the datasets of Cannon et al., 2016; Shackleton et al., 2019). Under similar ecological conditions, reference libraries could also be adapted to regional specificities, e.g. taxonomic differences between the Palearctic and Nearctic floras (Kuzmina et al., 2017).

The site map (Figure 1) highlights the nearly exclusive distribution of old (> 11,700 cal BP, i.e. Pleistocene) records in the Arctic, especially Siberia (e.g. meta-analysis by Wang et al., 2021). Indeed, current research suggests that a colder environment leads to a better preservation of “captive” eDNA, i.e. trapped in soil and sediment (Parducci et al., 2017; Taberlet et al., 2018). North America already has a comprehensive long-term macrofossil record for aquatic plants at the regional scale (Sawada et al., 2003), which foreshadows further reconstructions of freshwater postglacial conditions for that continent. However, the few existing studies for warmer regions, including one Eemian Ethiopian palaeolake, plus four Holocene records from Benin, Rwanda, and Namibia (Supplementary Table S2), show the potential for tropical regions. Thus, the likely success of such studies depends both on the completeness of a regional reference library adapted for the method, and the state of eDNA preservation, especially for ancient material.

Although we focused on freshwater ecosystems, Gaffney et al. (2020) also detected marine macrophytes (i.e. seagrasses), which represent entire families, e.g. Zosteraceae and Cymodoceaceae. Seagrasses play an important role in marine ecology, comparable in magnitude with coral reefs. Often growing in clonal colonies, they form large coastal habitats of paramount importance for the feeding and breeding of marine life, act as a carbon sink, and are a protection against coastal erosion (Arnaud-Haon et al., 2012; Christianen et al., 2013; Jiang et al., 2020; Lyimo, 2016; Zou et al., 2021). Seagrasses are well-detected in coastal sedimentary DNA (Foster et al., 2021; Ortega et al., 2020), and therefore their palaeorecord could also be used to reconstruct shallow marine environments in a similar way to our study. They may also be found in coastal lacustrine and fluvial environments, especially where there are large tidal ranges such as macrotidal estuaries.

### 4.2 Taxonomic resolution of the different methodological approaches

Our review covers three molecular approaches for biodiversity detection, the use of which depends on the aim of the study as these methodologies have different advantages and disadvantages (Box 1). Metagenomics and targeted capture are underrepresented in our dataset, as they have only recently been applied to eDNA and *sed*aDNA and therefore few studies have used these approaches for vascular plants, which is a limiting factor for the comparison of their respective taxonomic resolutions in aquatic macrophytes. To our knowledge, no direct comparison has yet been made between metabarcoding, metagenomics, and targeted capture to investigate yields with their respective optimal settings. Although Murchie et al. (2020) used all three methods, they used reference libraries whose respective taxonomic and biogeographic breadths differed by orders of magnitude: most of the Canadian flora is covered by at least one of three barcodes (*rbcL*, *matK*, ITS 2; Kuzmina et al., 2017) which is ideal for metabarcoding and targeted capture, but the more incomplete and uneven coverage for metagenomics hampered identification for the shotgun dataset. The best way to compare methods would be to analyse the same samples with the optimised pipeline of each, while using adequate and complete reference libraries.

Modern eDNA metabarcoding studies have used a range of markers, mainly long ones (usually > 400 bp); whereas the P6 loop is preferred in ancient studies. Indeed, the P6 loop is a short (51–135 bp) marker for vascular plants, and is thus especially suitable for degraded DNA (Taberlet et al., 2007). However, standard, longer (several hundreds of base pairs) barcodes are available for many more regions than P6 loop data, which provide opportunities for analyses of modern samples not limited by barcode length from more areas. Indeed, barcode taxonomic resolution usually increases with marker length (Figure 3). Most metabarcoding studies have used a single marker, but the method has potential for improvement by targeting multiple loci. Multi-marker studies often have used a combination of a nuclear and a chloroplast barcode: Kuzmina et al. (2018) used ITS 2 and *atp*B-*rbc*L; while Shackleton et al. (2019) combined 18S with the *trn*L P6 loop. All these markers were chosen while targeting a broader group than aquatic macrophytes, but it may be possible to develop new markers specifically for aquatic macrophytes, which have improved taxonomic resolution (Scriver et al., 2015). Nevertheless, even when targeting aquatics only, some loci fail to distinguish closely related species (e.g. within the Potamogetonaceae) because intraspecific and interspecific variability are of comparable magnitude (Du et al., 2011).

Shotgun sequencing is considered a less biased approach than metabarcoding for estimating abundance because it omits the PCR step which commonly causes biases (Pedersen et al., 2016). Metabarcoding and shotgun sequencing have both been used for nearly two decades (e.g. Willerslev et al., 2003; Tringe & Rubin, 2005), but the latter has greater sequencing costs. Most importantly, its efficiency relies on the availability of a considerably more complete reference library than a metabarcoding one, with entries from more target organisms and a broader genome coverage. As of today, genome skims are quite recent, and only available for the Arctic, part of the Chinese flora, and Western Australia (Alsos et al., 2020a; H.-T. Li et al., 2019; Nevill et al., 2020). Even genome skims cause unequal representation of taxa, and random match to closely related species may limit taxonomic resolution to genus level (Wang et al., 2021). While full genome assemblies are not available for any regional flora, the global number approaches 1,000 (Marks et al., 2021) and with time more will become available, enabling us to take full advantage of metagenomics. Yet, currently shotgun sequencing has a poor rate of raw sequence assignment compared to other methods: as its sequences originate from across entire genomes and any taxa and most of them do not reliably correspond to any known genomic reference (Peabody et al., 2015). For example most of the remainder are assigned to higher taxonomic ranks (e.g. kingdom, order), and only a minority gets identified to the genus or species level. Courtin et al. (2022) could only assign 12% of their shotgun sequencing reads, and Viridiplantae only accounted for 0.4% of these. This method requires comprehensive reference libraries, ideally entire genomes, as an incomplete reference library will cause many false positive matches as well as false negative misses.

The targeted capture approach consists of building a broad library of probes based on several hundreds to thousands of species, in order to cover the expected taxonomic breadth of broad groups such as vascular plants. It greatly increases the targeted region, at the cost of a diminished taxonomic resolution: Murchie et al. (2020) were not able to go much beyond genus level; and Krueger et al. (2021) only reported 6 macrophyte species and 1 genus, but did not report assignments at higher taxonomic levels. While it is in theory possible to identify more detected taxa to the species level, in practice most captured *sed*aDNA fragments are too short to be taxonomically informative at the species level, and DNA damage can lead to both false positives and negatives. However, detection of broad communities using targeted capture may have more promise for modern eDNA.

Our case study has a taxonomic resolution above average, with 68% of taxa identified to species level. However, some sequences remained unresolved between several species, sometimes even genera (e.g. *Nuphar* spp./*Nymphaea alba*). Because taxonomically unresolved sequences cannot always be assigned finite traits values, it in turn restricts the realised accuracy of our reconstruction. Taxonomic resolution in some aquatic taxa may be hampered by their close phylogeny, for instance within Potamogetonaceae. In our case study dataset, there were 8 sequences matching this family represented by 1,316,980 reads and 2,082 PCR replicates in total. The P6 loop sequence of Potamogetonaceae is longer than average (86 bp), and so the high number of reads shows that even longer sequences can be well amplified, which is probably due to the fact that aquatic macrophyte eDNA is better detected because they grow and decay adjacent to the sediment. There are eight *Potamogeton* and three *Stuckenia* species native to northern Fennoscandia (Alsos et al., 2022; Elven et al., 2022). *Potamogeton praelongus* has intraspecific variability, and different haplotypes may explain why our metabarcoding data did not match to PhyloNorway even if it did match at 100% with the three other reference libraries. Hybridisation, which is common in the genus *Potamogeton*, likely explains this sequence variability that may complicate taxonomic assignment both in reference libraries and consequently in metabarcoding data. Indeed, that metabarcoding dataset contains intraspecific variations as well as interspecific sequence sharing, but also one homopolymer consisting of six adenine bases, which can cause problems during amplification and sequencing. For studies targeting these taxa alternative barcodes should be explored.

At the moment, metabarcoding stands out as the method with the highest taxonomic resolution, both for aquatic macrophytes and for the total flora. Recent advances have demonstrated that it is possible to assemble genome-scale reference libraries, including from herbarium collections (Alsos et al., 2020a). This can assist in unlocking the bottleneck to access the large potential of metagenomics for palaeoecology of aquatic macrophytes and other taxa (Wang et al. 2021; Kjær et al. 2022). We can expect to see a continued increase in eDNA publications in the near future (Capo et al., 2021), as methods are refined and reference libraries are supplemented. Metabarcoding is the least expensive of all three methods and requires smaller reference libraries. It is also analytically easier, as the genomic locus is constrained and full barcode sequences are generated, unlike either of the metagenomic-based methods; thus it might remain popular for modern surveys of aquatic macrophyte diversity in the short-to-medium term. For palaeoecological studies, it is crucial to use a short marker in order to capture the diversity of taxa represented in ancient material, and in turn provide an overview of best-estimate of past communities.

### 4.3 Potential for ecological and environmental reconstruction

Environmental traits have different temporal trends: nitrogen availability appears to remain stable throughout the Holocene, while light optimum changes gradually and temperature shifts are more abrupt. We have reconstructed the continuous postglacial ecological history of northern Fennoscandia. This gives a general overview of regional conditions, and follows the Fennoscandian ice sheet retreat as lakes become ice-free (Alsos et al., 2022). Here, the ecological succession of aquatic communities retraces the establishment of modern environmental conditions since the onset of the Holocene. Aquatic macrophytes appear later in lake records than any other plant functional group, but contrary to the terrestrial vegetation, their diversity soars rapidly after arrival (Alsos et al., 2022).

Environmental trends are also visible among lakes, and these site-specific conditions emphasise the local scale precision achieved by our reconstruction, precisely because aquatic macrophytes have restricted niches related to water level. The remarkable environmental stability in Sandfjorddalen throughout the Holocene demonstrates that niche persistence over time existed even in the early record. Conversely, other lakes displayed fluctuating conditions, offering evolving niches for aquatic plant communities. This suggests that even their early occurrences can be used confidently for environmental reconstruction. Indeed, the temporal compositional and richness changes could be observed across Siberia and Tibet for the Pleistocene and Holocene (Stoof-Leichsenring et al., 2022). Combining the modern dataset with the core metabarcoding data would allow to use the macrophyte community signal for the reconstruction of past environments, such as conductivity and temperature.

Beyond ecological and environmental reconstructions, the *sed*aDNA signal from macrophytes could in the future be leveraged to address evolutionary questions. High-resolution time series can be combined with other proxies in order to allow statistical analyses and inference of causal relationships among drivers of ecosystem changes, such as human land use and climate change (Garcés-Pastor et al., 2022). With the rapid ongoing methodological improvements, distinguishing closely related species and even haplotypes could support the fine-scale mapping of genetic diversity and its variation in time and space (Epp et al., 2018), and help retrace postglacial colonisation routes to solve the origin of modern boreal and arctic vegetation. Lastly, the recent retrieval of geological age eDNA (Kjær et al., 2022) could give a direct insight on actual genetic evolutive processes at the million-years timescale, especially for organisms such as macrophytes which rarely persist as macrofossils.

In sum, macrophyte-based P6 loop metabarcoding supports reconstructions for a wide array of important freshwater metrics. From *sed*aDNA-derived macrophyte turnover and community composition, we can infer lake level changes (Heinecke et al., 2017) and conductivity and temperature changes (Stoof-Leichsenring et al., 2022) on millennial time scales. Our study extends environmental reconstructions for northern Fennoscandian lakes over the last twelve thousand years to thermal range, continentality, nutrient availability, light conditions, and water pH. We show the potential that aquatic plants aDNA offers to reconstruct past environments when using a comprehensive trait database of the local flora.

## 5. Conclusion

This review reveals that aquatic macrophytes are often detected in both eDNA and *sed*aDNA studies, being often the most common and abundant taxa. Environmental DNA from aquatics is reported in a wide range of sample types, from stream water to permafrost, and in various bioregions although colder regions are overrepresented at present. *Sed*aDNA of aquatic macrophytes can be detected from samples over two millions years old in favourable environments (Kjær et al. 2022), which indicates a considerable potential for long-term *sed*aDNA-based environmental reconstruction.

Shotgun sequencing appears to lack sufficient taxonomic resolution to identify environmentally indicative aquatic macrophyte taxa at present. Targeted capture lacks more application on plants, but aDNA results on mammals are encouraging for the development of bait-sets that could screen past and present plant biodiversity. Barcoding can track a single species through space, working under the assumption of presence of the targeted taxon; yet its potential when applied to aDNA is limited by the length of barcode used. Metabarcoding is a more versatile tool, and there is a wide range of markers one can choose from. However, when applying this approach to ancient samples, we recommend the use of a short (< ca. 100 bp) marker in order to maximise the yield, but still able to capture and identify sequences assuming the availability of a corresponding reference library. In this regard, the *g*–*h* primer set targeting the P6 loop fulfils these requirements to a large extent and it has proven successful in a variety of studies, due to the availability of highly curated sequence libraries.

Using *sed*aDNA to reconstruct ecological and environmental conditions of the past can be supplemented by cross-disciplinary expertise on local and regional diversity, postglacial migration, vegetation cover and land use changes (Alsos et al., 2022; Brown et al., 2022). The paramount role of reference libraries is highlighted by the differences in taxonomic resolution between methods as the requirements for targeted capture largely vary depending on the taxonomic diversity targeted; metabarcoding often goes to the species level due to well-curated barcode databases; while metagenomics could potentially achieve comparable results but that will require increased sequencing reference genomes.

## Author contributions

Conceptualisation: IGA. Conducting the research: AR, PDH, AGB, DPR, IGA. Data analysis: AR, DPR. Preparation of figures & tables: AR. Data interpretation, writing: AR, DPR, PDH, AGB, KSL, IGA.

## Data availability

Full references for the reviewed papers, and data extracted from it, are found in Supplementary Table S1 (by publication) and Supplementary Table S2 (by site). The full list of aquatic taxa detected is given as Supplementary Table S3. The *sed*aDNA dataset from Rijal et al. (2021) can be found in their supplementary information; the reanalysed version used here can be found in Alsos et al. (2022). The trait database is available as supplementary information in Tyler et al. (2021).

## Acknowledgements

This work was supported by the ECOGEN project (Ecosystem change and species persistence over time: a genome-based approach), Research Council of Norway grant 226134/F50 (to I.G.A.). We thank Pablo Raguet for his help with data analysis and visualisation.

## Supplementary information

- Table S1: List of the studies reviewed, references, and methodological information
- Table S2: List of sites corresponding to the reviewed studies
- Table S3: List of taxa detected by the reviewed studies and their classification as aquatic or not
- Table S4: Comparison of various aquatic plant classifications to the moisture trait from Tyler et al. (2021)
- Table S5: Wilcoxon tests for differences in taxonomic resolution between aquatics and all vascular taxa by method
- Figure S1: Plots of the Generalised Linear Models investigating how the year of publication and age of the oldest sample affect the total richness of aquatic macrophytes detected
- Figure S2: Correlogram of the preselected environmental traits
- Figure S3: Lake-specific richness in aquatic macrophyte taxa through time
- Figure S4: Graphical visualisation of environmental niches per taxon, for the selected traits
- Figure S5: Proportion of traits values over time in PCR replicates for lakes (a) Horntjernet, (b) Sandfjorddalen, (c) Kuutsjärvi, (d) Nordvivatnet, and (d) Sierravannet
- Figure S6: Proportion of PCR replicates for each taxon over time for lakes (a) Gauptjern, (b) Sandfjorddalen, (c) Horntjernet, (d) Kuutsjärvi, (e) Nordvivatnet, (f) Sierravannet, (g) Nesservatnet, (h) Eaštorjávri South, (i) Jøkelvatnet, and (j) Langfjordvannet

## References

Ackerfield, J. (2015). *Flora of Colorado* (Vol. 41). Botanical Research Institute of Texas. https://www.nhbs.com/flora-of-colorado-book

Adame, M. F., & Reef, R. (2020). Potential Pollution Sources from Agricultural Activities on Tropical Forested Floodplain Wetlands Revealed by Soil eDNA. Forests, 11(8), Article 8. https://doi.org/10.3390/f11080892

Alahuhta, J., Rosbakh, S., Chepinoga, V., & Heino, J. (2020). Environmental determinants of lake macrophyte communities in Baikal Siberia. Aquatic Sciences, 82(2), 39. https://doi.org/10.1007/s00027-020-0710-8

Alsos, I. G., Lammers, Y., Kjellman, S. E., Merkel, M. K. F., Bender, E. M., Rouillard, A., Erlendsson, E., Guðmundsdóttir, E. R., Benediktsson, Í. Ö., Farnsworth, W. R., Brynjólfsson, S., Gísladóttir, G., Eddudóttir, S. D., & Schomacker, A. (2021). Ancient sedimentary DNA shows rapid post-glacial colonisation of Iceland followed by relatively stable vegetation until the Norse settlement (Landnám) AD 870. Quaternary Science Reviews, 259, 106903. https://doi.org/10.1016/j.quascirev.2021.106903

Alsos, I. G., Lammers, Y., Yoccoz, N. G., Jørgensen, T., Sjögren, P., Gielly, L., & Edwards, M. E. (2018). Plant DNA metabarcoding of lake sediments: How does it represent the contemporary vegetation. PLOS ONE, 13(4), e0195403. https://doi.org/10.1371/journal.pone.0195403

Alsos, I. G., Rijal, D. P., Ehrich, D., Karger, D. N., Yoccoz, N. G., Heintzman, P. D., Brown, A. G., Lammers, Y., Pellissier, L., Alm, T., Bråthen, K. A., Coissac, E., Merkel, M. K. F., Alberti, A., Denoeud, F., Bakke, J., & PHYLONORWAY CONSORTIUM. (2022). Postglacial species arrival and diversity buildup of northern ecosystems took millennia. Science Advances, 8(39), eabo7434. https://doi.org/10.1126/sciadv.abo7434

Alsos, I. G., Sjögren, P., Brown, A. G., Gielly, L., Merkel, M. K. F., Paus, A., Lammers, Y., Edwards, M. E., Alm, T., Leng, M., Goslar, T., Langdon, C. T., Bakke, J., & van der Bilt, W. G. M. (2020b). Last Glacial Maximum environmental conditions at Andøya, northern Norway; evidence for a northern ice-edge ecological “hotspot.” Quaternary Science Reviews, 239, 106364. https://doi.org/10.1016/j.quascirev.2020.106364

Alsos, I. G., Sjögren, P., Edwards, M. E., Landvik, J. Y., Gielly, L., Forwick, M., Coissac, E., Brown, A. G., Jakobsen, L. V., Føreid, M. K., & Pedersen, M. W. (2016). Sedimentary ancient DNA from Lake Skartjørna, Svalbard: Assessing the resilience of arctic flora to Holocene climate change. The Holocene, 26(4), 627–642. https://doi.org/10.1177/0959683615612563

Anglès d’Auriac, M. B., Strand, D. A., Mjelde, M., Demars, B. O. L., & Thaulow, J. (2019). Detection of an invasive aquatic plant in natural water bodies using environmental DNA. PLOS ONE, 14(7), e0219700. https://doi.org/10.1371/journal.pone.0219700

Arnaud-Haond, S., Duarte, C. M., Diaz-Almela, E., Marbà, N., Sintes, T., & Serrão, E. A. (2012). Implications of Extreme Life Span in Clonal Organisms: Millenary Clones in Meadows of the Threatened Seagrass Posidonia oceanica. PLOS ONE, 7(2), e30454. https://doi.org/10.1371/journal.pone.0030454

Australian Biological Resources Study. (2015). Flora of Australia (Vol. 1–59). CSIRO Publishing.

Bjune, A. E., Greve Alsos, I., Brendryen, J., Edwards, M. E., Haflidason, H., Johansen, M. S., Mangerud, J., Paus, A., Regnéll, C., Svendsen, J.-I., & Clarke, C. L. (2021). Rapid climate changes during the Lateglacial and the early Holocene as seen from plant community dynamics in the Polar Urals, Russia. Journal of Quaternary Science, n/a(n/a). https://doi.org/10.1002/jqs.3352

Boessenkool, S., McGlynn, G., Epp, L. S., Taylor, D., Pimentel, M., Gizaw, A., Nemomissa, S., Brochmann, C., & Popp, M. (2014). Use of Ancient Sedimentary DNA as a Novel Conservation Tool for High-Altitude Tropical Biodiversity. Conservation Biology, 28(2), 446–455. https://doi.org/10.1111/cobi.12195

Bremond, L., Favier, C., Ficetola, G. F., Tossou, M. G., Akouégninou, A., Gielly, L., Giguet-Covex, C., Oslisly, R., & Salzmann, U. (2017). Five thousand years of tropical lake sediment DNA records from Benin. Quaternary Science Reviews, 170, 203–211. https://doi.org/10.1016/j.quascirev.2017.06.025

Brooks, S. J., & Birks, H. J. B. (2000). Chironomid-inferred late-glacial and early-Holocene mean July air temperatures for Kråkenes Lake, western Norway. Journal of Paleolimnology, 23(1), 77–89. https://doi.org/10.1023/A:1008044211484

Brown, A. G. (2002). Learning from the past: Palaeohydrology and palaeoecology. Freshwater Biology, 47(4), 817–829.

Brown, T., Rijal, D. P., Heintzman, P. D., Clarke, C. L., Blankholm, H.-P., Høeg, H. I., Lammers, Y., Bråthen, K. A., Edwards, M., & Alsos, I. G. (2022). Paleoeconomy more than demography determined prehistoric human impact in Arctic Norway. *PNAS Nexus*, pgac209. https://doi.org/10.1093/pnasnexus/pgac209

Brown, A. G., Van Hardenbroek, M., Fonville, T., Davies, K., Mackay, H., Murray, E., Head, K., Barratt, P., McCormick, F., Ficetola, G. F., Gielly, L., Henderson, A. C. G., Crone, A., Cavers, G., Langdon, P. G., Whitehouse, N. J., Pirrie, D., & Alsos, I. G. (2021). Ancient DNA, lipid biomarkers and palaeoecological evidence reveals construction and life on early medieval lake settlements. Scientific Reports, 11(1), 11807. https://doi.org/10.1038/s41598-021-91057-x

Buxton, A. S., Groombridge, J. J., & Griffiths, R. A. (2018). Seasonal variation in environmental DNA detection in sediment and water samples. PLOS ONE, 13(1), e0191737. https://doi.org/10.1371/journal.pone.0191737

Cannon, M. V., Hester, J., Shalkhauser, A., Chan, E. R., Logue, K., Small, S. T., & Serre, D. (2016). In silico assessment of primers for eDNA studies using PrimerTree and application to characterize the biodiversity surrounding the Cuyahoga River. Scientific Reports, 6(1), Article 1. https://doi.org/10.1038/srep22908

Capo, E., Giguet-Covex, C., Rouillard, A., Nota, K., Heintzman, P. D., Vuillemin, A., Ariztegui, D., Arnaud, F., Belle, S., Bertilsson, S., Bigler, C., Bindler, R., Brown, A. G., Clarke, C. L., Crump, S. E., Debroas, D., Englund, G., Ficetola, G. F., Garner, R. E., … Parducci, L. (2021). Lake Sedimentary DNA Research on Past Terrestrial and Aquatic Biodiversity: Overview and Recommendations. Quaternary, 4(1), 6. https://doi.org/10.3390/quat4010006

Christianen, M. J. A., Belzen, J. van, Herman, P. M. J., Katwijk, M. M. van, Lamers, L. P. M., Leent, P. J. M. van, & Bouma, T. J. (2013). Low-Canopy Seagrass Beds Still Provide Important Coastal Protection Services. PLOS ONE, 8(5), e62413. https://doi.org/10.1371/journal.pone.0062413

Clarke, C. L., Alsos, I. G., Edwards, M. E., Paus, A., Gielly, L., Haflidason, H., Mangerud, J., Regnéll, C., Hughes, P. D. M., Svendsen, J. I., & Bjune, A. E. (2020). A 24,000-year ancient DNA and pollen record from the Polar Urals reveals temporal dynamics of arctic and boreal plant communities. Quaternary Science Reviews, 247, 106564. https://doi.org/10.1016/j.quascirev.2020.106564

Clarke, C. L., Edwards, M. E., Brown, A. G., Gielly, L., Lammers, Y., Heintzman, P. D., Ancin-Murguzur, F. J., Bråthen, K.-A., Goslar, T., & Alsos, I. G. (2019). Holocene floristic diversity and richness in northeast Norway revealed by sedimentary ancient DNA (sedaDNA) and pollen. Boreas, 48(2), 299–316. https://doi.org/10.1111/bor.12357

Clayton, W. D., Phillips, S. M., & Renvoize, S. A. (1974). Flora of Tropical East Africa. Gramineae (Part 2). Royal Botanic Gardens, Kew. https://www.cabdirect.org/cabdirect/abstract/19740724436

Coghlan, S. A., Shafer, A. B. A., & Freeland, J. R. (2021). Development of an environmental DNA metabarcoding assay for aquatic vascular plant communities. Environmental DNA, 3(2), 372–387. https://doi.org/10.1002/edn3.120

Courtin, J., Andreev, A. A., Raschke, E., Bala, S., Biskaborn, B. K., Liu, S., Zimmermann, H., Diekmann, B., Stoof-Leichsenring, K. R., Pestryakova, L. A., & Herzschuh, U. (2021). Vegetation Changes in Southeastern Siberia During the Late Pleistocene and the Holocene. Frontiers in Ecology and Evolution, 9, 18.

Courtin, J., Perfumo, A., Andreev, A. A., Opel, T., Stoof-Leichsenring, K. R., Edwards, M. E., Murton, J. B., & Herzschuh, U. (2022). Pleistocene glacial and interglacial ecosystems inferred from ancient DNA analyses of permafrost sediments from Batagay megaslump, East Siberia. Environmental DNA, 4(6), 1265–1283. https://doi.org/10.1002/edn3.336

Crump, S. E., Fréchette, B., Power, M., Cutler, S., de Wet, G., Raynolds, M. K., Raberg, J. H., Briner, J. P., Thomas, E. K., Sepúlveda, J., Shapiro, B., Bunce, M., & Miller, G. H. (2021). Ancient plant DNA reveals High Arctic greening during the Last Interglacial. Proceedings of the National Academy of Sciences, 118(13). https://doi.org/10.1073/pnas.2019069118

Crump, S. E., Miller, G. H., Power, M., Sepúlveda, J., Dildar, N., Coghlan, M., & Bunce, M. (2019). Arctic shrub colonization lagged peak postglacial warmth: Molecular evidence in lake sediment from Arctic Canada. Global Change Biology, 25(12), 4244–4256. https://doi.org/10.1111/gcb.14836

Curtin, L., D’Andrea, W. J., Balascio, N. L., Shirazi, S., Shapiro, B., de Wet, G. A., Bradley, R. S., & Bakke, J. (2021). Sedimentary DNA and molecular evidence for early human occupation of the Faroe Islands. Communications Earth & Environment, 2(1), Article 1. https://doi.org/10.1038/s43247-021-00318-0

Dalla Vecchia, A., Villa, P., & Bolpagni, R. (2020). Functional traits in macrophyte studies: Current trends and future research agenda. Aquatic Botany, 167, 103290. https://doi.org/10.1016/j.aquabot.2020.103290

Dan, Z., Chuan, W., Qiaohong, Z., & Xingzhong, Y. (2021). Sediments nitrogen cycling influenced by submerged macrophytes growing in winter. Water Science and Technology, 83(7), 1728–1738. https://doi.org/10.2166/wst.2021.081

Dar, N. A., Pandit, A. K., & Ganai, B. A. (2014). Factors affecting the distribution patterns of aquatic macrophytes. Limnological Review, 14(2), 75–81. https://doi.org/10.2478/limre-2014-0008

Doi, H., Akamatsu, Y., Goto, M., Inui, R., Komuro, T., Nagano, M., & Minamoto, T. (2021). Broad-scale detection of environmental DNA for an invasive macrophyte and the relationship between DNA concentration and coverage in rivers. Biological Invasions, 23(2), 507–520. https://doi.org/10.1007/s10530-020-02380-9

Drummond, J. A., Larson, E. R., Li, Y., Lodge, D. M., Gantz, C. A., Pfrender, M. E., Renshaw, M. A., Correa, A. M. S., & Egan, S. P. (2021). Diversity Metrics Are Robust to Differences in Sampling Location and Depth for Environmental DNA of Plants in Small Temperate Lakes. Frontiers in Environmental Science, 9. https://www.frontiersin.org/article/10.3389/fenvs.2021.617924

Du, Z.-Y., Qimike, A., Yang, C.-F., Chen, J.-M., & Wang, Q.-F. (2011). Testing four barcoding markers for species identification of Potamogetonaceae. Journal of Systematics and Evolution, 49(3), 246–251. https://doi.org/10.1111/j.1759-6831.2011.00131.x

Ellenberg, H., Weber, H. E., Düll, R., Wirth, V., & Werner, W. (2001). Zeigerwerte von Pflanzen in Mitteleuropa. Scripta Geobotanica, 18.

Elven, R., Arnesen, G., Alsos, I. G., & Sandbakk, B. (2020). *SvalbardFlora*. https://svalbardflora.no

Elven, R., Sletten Bjorå, C., Fremstad, E., Hegre, H., & Solstad, H. (2022). *Norsk Flora* (8th ed.). Det Norske Samlaget. https://www.naturogfritid.no/860109/Norsk+Flora

Epp, L. S., Kruse, S., Kath, N. J., Stoof-Leichsenring, K. R., Tiedemann, R., Pestryakova, L. A., & Herzschuh, U. (2018). Temporal and spatial patterns of mitochondrial haplotype and species distributions in Siberian larches inferred from ancient environmental DNA and modeling. Scientific Reports, 8(1), 17436. https://doi.org/10.1038/s41598-018-35550-w

Fellows Yates, J. A., Andrades Valtueña, A., Vågene, Å. J., Cribdon, B., Velsko, I. M., Borry, M., Bravo-Lopez, M. J., Fernandez-Guerra, A., Green, E. J., Ramachandran, S. L., Heintzman, P. D., Spyrou, M. A., Hübner, A., Gancz, A. S., Hider, J., Allshouse, A. F., Zaro, V., & Warinner, C. (2021). Community-curated and standardised metadata of published ancient metagenomic samples with AncientMetagenomeDir. Scientific Data, 8(1), 31. https://doi.org/10.1038/s41597-021-00816-y

Foster, N. R., van Dijk, K., Biffin, E., Young, J. M., Thomson, V. A., Gillanders, B. M., Jones, A. R., & Waycott, M. (2021). A Multi-Gene Region Targeted Capture Approach to Detect Plant DNA in Environmental Samples: A Case Study From Coastal Environments. Frontiers in Ecology and Evolution, 9. https://www.frontiersin.org/article/10.3389/fevo.2021.735744

Fujiwara, A., Matsuhashi, S., Doi, H., Yamamoto, S., & Minamoto, T. (2016). Use of environmental DNA to survey the distribution of an invasive submerged plant in ponds. Freshwater Science, 35(2), 748–754. https://doi.org/10.1086/685882

Gaffney, V., Fitch, S., Bates, M., Ware, R. L., Kinnaird, T., Gearey, B., Hill, T., Telford, R., Batt, C., Stern, B., Whittaker, J., Davies, S., Sharada, M. B., Everett, R., Cribdon, R., Kistler, L., Harris, S., Kearney, K., Walker, J., … Allaby, R. G. (2020). Multi-Proxy Characterisation of the Storegga Tsunami and Its Impact on the Early Holocene Landscapes of the Southern North Sea. Geosciences, 10(7), 270. https://doi.org/10.3390/geosciences10070270

Gantz, C. A., Renshaw, M. A., Erickson, D., Lodge, D. M., & Egan, S. P. (2018). Environmental DNA detection of aquatic invasive plants in lab mesocosm and natural field conditions. Biological Invasions, 20(9), 2535–2552. https://doi.org/10.1007/s10530-018-1718-z

Garcés-Pastor, S., Coissac, E., Lavergne, S., Schwörer, C., Theurillat, J.-P., Heintzman, P. D., Wangensteen, O. S., Tinner, W., Rey, F., Heer, M., Rutzer, A., Walsh, K., Lammers, Y., Brown, A. G., Goslar, T., Rijal, D. P., Karger, D. N., Pellissier, L., Heiri, O., & Alsos, I. G. (2022). High resolution ancient sedimentary DNA shows that alpine plant diversity is associated with human land use and climate change. Nature Communications, 13(1), Article 1. https://doi.org/10.1038/s41467-022-34010-4

GBIF.org. (2022). *Global Biodiversity Information Facility*. https://www.gbif.org

Giguet-Covex, C., Ficetola, G. F., Walsh, K., Poulenard, J., Bajard, M., Fouinat, L., Sabatier, P., Gielly, L., Messager, E., Develle, A. L., David, F., Taberlet, P., Brisset, E., Guiter, F., Sinet, R., & Arnaud, F. (2019). New insights on lake sediment DNA from the catchment: Importance of taphonomic and analytical issues on the record quality. Scientific Reports, 9, 21.

Haines, A., Farnsworth, E. J., Morrison, G., & New England Wild Flower Society. (2011). New England Wild Flower Society’s Flora Novae Angliae A Manual for the Identification of Native and Naturalized Higher Vascular Plants of New England. Yale University Press.

Harper, L. R., Buxton, A. S., Rees, H. C., Bruce, K., Brys, R., Halfmaerten, D., Read, D. S., Watson, H. V., Sayer, C. D., Jones, E. P., Priestley, V., Mächler, E., Múrria, C., Garcés-Pastor, S., Medupin, C., Burgess, K., Benson, G., Boonham, N., Griffiths, R. A., … Hänfling, B. (2019). Prospects and challenges of environmental DNA (eDNA) monitoring in freshwater ponds. Hydrobiologia, 826(1), 25–41. https://doi.org/10.1007/s10750-018-3750-5

Heinecke, L., Epp, L. S., Reschke, M., Stoof-Leichsenring, K. R., Mischke, S., Plessen, B., & Herzschuh, U. (2017). Aquatic macrophyte dynamics in Lake Karakul (Eastern Pamir) over the last 29 cal ka revealed by sedimentary ancient DNA and geochemical analyses of macrofossil remains. Journal of Paleolimnology, 58(3), 403–417. https://doi.org/10.1007/s10933-017-9986-7

Hollingsworth, P. M., Graham, S. W., & Little, D. P. (2011). Choosing and Using a Plant DNA Barcode. PLOS ONE, 6(5), e19254. https://doi.org/10.1371/journal.pone.0019254

Huang, S., Stoof-Leichsenring, K. R., Liu, S., Courtin, J., Andreev, A. A., Pestryakova, Luidmila. A., & Herzschuh, U. (2021). Plant Sedimentary Ancient DNA From Far East Russia Covering the Last 28,000 Years Reveals Different Assembly Rules in Cold and Warm Climates. Frontiers in Ecology and Evolution, 9, 873. https://doi.org/10.3389/fevo.2021.763747

Hughes, A. L. C., Gyllencreutz, R., Lohne, Ø. S., Mangerud, J., & Svendsen, J. I. (2016). The last Eurasian ice sheets – a chronological database and time-slice reconstruction, DATED-1. Boreas, 45(1), 1–45. https://doi.org/10.1111/bor.12142

Hughes, J. M. R., Clarkson, B. R., Castro-Castellon, A. T., & Hess, L. L. (2018). Wetland Plants and Aquatic Macrophytes. In J. M. R. Hughes (Ed.), Freshwater Ecology and Conservation: Approaches and Techniques (p. 480). Oxford University Press.

Ibrahim, A., Höckendorff, S., Schleheck, D., Epp, L., van Kleunen, M., & Meyer, A. (2022). Vegetation changes over the last centuries in the Lower Lake Constance region reconstructed from sediment-core environmental DNA. Environmental DNA, 4(4), 830–845. https://doi.org/10.1002/edn3.292

Info Flora. (2022). *Info Flora*. Info Flora CH. https://www.infoflora.ch/en/

Ji, F., Yan, L., Yan, S., Qin, T., Shen, J., & Zha, J. (2021). Estimating aquatic plant diversity and distribution in rivers from Jingjinji region, China, using environmental DNA metabarcoding and a traditional survey method. Environmental Research, 199, 111348. https://doi.org/10.1016/j.envres.2021.111348

Jia, W., Liu, X., Stoof-Leichsenring, K. R., Liu, S., Li, K., & Herzschuh, U. (2021). Preservation of sedimentary plant DNA is related to lake water chemistry. Environmental DNA, 4(2). https://doi.org/10.1002/edn3.259

Jiang, Z., Huang, D., Fang, Y., Cui, L., Zhao, C., Liu, S., Wu, Y., Chen, Q., Ranvilage, C. I. P. M., He, J., & Huang, X. (2020). Home for Marine Species: Seagrass Leaves as Vital Spawning Grounds and Food Source. Frontiers in Marine Science, 7. https://www.frontiersin.org/articles/10.3389/fmars.2020.00194

Johnson, R. K., & Toprak, V. (2021). Local habitat is a strong determinant of spatial and temporal patterns of macrophyte diversity and composition in boreal lakes. Freshwater Biology, 66(8), 1490–1501. https://doi.org/10.1111/fwb.13733

Jørgensen, T., Haile, J., Möller, P., Andreev, A., Boessenkool, S., Rasmussen, M., Kienast, F., Coissac, E., Taberlet, P., Brochmann, C., Bigelow, N. H., Andersen, K., Orlando, L., Gilbert, M. T. P., & Willerslev, E. (2012). A comparative study of ancient sedimentary DNA, pollen and macrofossils from permafrost sediments of northern Siberia reveals long-term vegetational stability. Molecular Ecology, 21(8), 1989–2003. https://doi.org/10.1111/j.1365-294X.2011.05287.x

Kisand, V., Talas, L., Kisand, A., Stivrins, N., Reitalu, T., Alliksaar, T., Vassiljev, J., Liiv, M., Heinsalu, A., Seppä, H., & Veski, S. (2018). From microbial eukaryotes to metazoan vertebrates: Wide spectrum paleo- diversity in sedimentary ancient DNA over the last 14,500 years. Geobiology, 16(6), 628–639. https://doi.org/10.1111/gbi.12307

Kjær, K. H., Winther Pedersen, M., De Sanctis, B., De Cahsan, B., Korneliussen, T. S., Michelsen, C. S., Sand, K. K., Jelavić , S., Ruter, A. H., Schmidt, A. M. A., Kjeldsen, K. K., Tesakov, A. S., Snowball, I., Gosse, J. C., Alsos, I. G., ang, Y., Dockter, C., Rasmussen, M., Jørgensen, M. E., … Willerslev, E. (2022). A 2-million-year-old ecosystem in Greenland uncovered by environmental DNA. Nature, 612(7939), Article 7939. https://doi.org/10.1038/s41586-022-05453-y

Kodama, T., Miyazono, S., Akamatsu, Y., Tsuji, S., & Nakao, R. (2022). Abundance estimation of riverine macrophyte Egeria densa using environmental DNA: Effects of sampling season and location. Limnology, 23(2), 299–308. https://doi.org/10.1007/s10201-021-00689-5

Krueger, J., Foerster, V., Trauth, M. H., Hofreiter, M., & Tiedemann, R. (2021). Exploring the Past Biosphere of Chew Bahir/Southern Ethiopia: Cross-Species Hybridization Capture of Ancient Sedimentary DNA from a Deep Drill Core. Frontiers in Earth Science, 9. https://www.frontiersin.org/articles/10.3389/feart.2021.683010

Kuehne, L. M., Ostberg, C. O., Chase, D. M., Duda, J. J., & Olden, J. D. (2020). Use of environmental DNA to detect the invasive aquatic plants Myriophyllum spicatum and Egeria densa in lakes. Freshwater Science, 39(3), 521–533. https://doi.org/10.1086/710106

Kuzmina, M. L., Braukmann, T. W. A., Fazekas, A. J., Graham, S. W., Dewaard, S. L., Rodrigues, A., Bennett, B. A., Dickinson, T. A., Saarela, J. M., Catling, P. M., Newmaster, S. G., Percy, D. M., Fenneman, E., Lauron-Moreau, A., Ford, B., Gillespie, L., Subramanyam, R., Whitton, J., Jennings, L., … Hebert, P. D. N. (2017). Using Herbarium-Derived DNAs to Assemble a Large-Scale DNA Barcode Library for the Vascular Plants of Canada. Applications in Plant Sciences, 5(12), 1700079. https://doi.org/10.3732/apps.1700079

Kuzmina, M. L., Braukmann, T. W. A., & Zakharov, E. V. (2018). Finding the pond through the weeds: EDNA reveals underestimated diversity of pondweeds. Applications in Plant Sciences, 6(5), e01155. https://doi.org/10.1002/aps3.1155

Li, H.-T., Yi, T.-S., Gao, L.-M., Ma, P.-F., Zhang, T., Yang, J.-B., Gitzendanner, M. A., Fritsch, P. W., Cai, J., Luo, Y., Wang, H., van der Bank, M., Zhang, S.-D., Wang, Q.-F., Wang, J., Zhang, Z.-R., Fu, C.-N., Yang, J., Hollingsworth, P. M., … Li, D.-Z. (2019). Origin of angiosperms and the puzzle of the Jurassic gap. Nature Plants, 5(5), 461–470. https://doi.org/10.1038/s41477-019-0421-0

Li, H., Zhang, H., Chang, F., Liu, Q., Zhang, Y., Liu, F., & Zhang, X. (2023). Sedimentary DNA for tracking the long-term changes in biodiversity. Environmental Science and Pollution Research. https://doi.org/10.1007/s11356-023-25130-5

Liu, S., Kruse, S., Scherler, D., Ree, R. H., Zimmermann, H. H., Stoof-Leichsenring, K. R., Epp, L. S., Mischke, S., & Herzschuh, U. (2021). Sedimentary ancient DNA reveals a threat of warming-induced alpine habitat loss to Tibetan Plateau plant diversity. Nature Communications, 12(1), 2995. https://doi.org/10.1038/s41467-021-22986-4

Liu, S., Stoof-Leichsenring, K. R., Kruse, S., Pestryakova, L. A., & Herzschuh, U. (2020). Holocene Vegetation and Plant Diversity Changes in the North-Eastern Siberian Treeline Region From Pollen and Sedimentary Ancient DNA. Frontiers in Ecology and Evolution, 8, 304. https://doi.org/10.3389/fevo.2020.560243

Lyimo, L. D. (2016). Carbon sequestration processes in tropical seagrass beds [Stockholm University, Faculty of Science, Department of Ecology, Environment and Plant Sciences. University of Dodoma.]. http://urn.kb.se/resolve?urn=urn:nbn:se:su:diva-128201

Maasri, A., Jähnig, S. C., Adamescu, M. C., Adrian, R., Baigun, C., Baird, D. J., Batista-Morales, A., Bonada, N., Brown, L. E., Cai, Q., Campos-Silva, J. V., Clausnitzer, V., Contreras-MacBeath, T., Cooke, S. J., Datry, T., Delacámara, G., De Meester, L., Dijkstra, K.-D. B., Do, V. T., … Worischka, S. (2022). A global agenda for advancing freshwater biodiversity research. Ecology Letters, 25(2), 255–263. https://doi.org/10.1111/ele.13931

Maddison, D. R., & Schulz, K.-S. (2007). *The Tree of Life Web Project*. The Tree of Life Web Project. http://tolweb.org

Marks, R. A., Hotaling, S., Frandsen, P. B., & VanBuren, R. (2021). Representation and participation across 20 years of plant genome sequencing. Nature Plants, 7(12), Article 12. https://doi.org/10.1038/s41477-021-01031-8

Matsuhashi, S., Doi, H., Fujiwara, A., Watanabe, S., & Minamoto, T. (2016). Evaluation of the Environmental DNA Method for Estimating Distribution and Biomass of Submerged Aquatic Plants. PLOS ONE, 11(6), e0156217. https://doi.org/10.1371/journal.pone.0156217

Murchie, T. J., Kuch, M., Duggan, A. T., Ledger, M. L., Roche, K., Klunk, J., Karpinski, E., Hackenberger, D., Sadoway, T., MacPhee, R., Froese, D., & Poinar, H. (2020). Optimizing extraction and targeted capture of ancient environmental DNA for reconstructing past environments using the PalaeoChip Arctic-1.0 bait-set. Quaternary Research, 99, 305–328. https://doi.org/10.1017/qua.2020.59

Murphy, K., Efremov, A., Davidson, T. A., Molina-Navarro, E., Fidanza, K., Crivelari Betiol, T. C., Chambers, P., Tapia Grimaldo, J., Varandas Martins, S., Springuel, I., Kennedy, M., Mormul, R. P., Dibble, E., Hofstra, D., Lukács, B. A., Gebler, D., Baastrup-Spohr, L., & Urrutia-Estrada, J. (2019). World distribution, diversity and endemism of aquatic macrophytes. Aquatic Botany, 158, 103127. https://doi.org/10.1016/j.aquabot.2019.06.006

Nevill, P. G., Zhong, X., Tonti-Filippini, J., Byrne, M., Hislop, M., Thiele, K., van Leeuwen, S., Boykin, L. M., & Small, I. (2020). Large scale genome skimming from herbarium material for accurate plant identification and phylogenomics. Plant Methods, 16(1), 1. https://doi.org/10.1186/s13007-019-0534-5

Newton, J., Sepulveda, A., Sylvester, K., & Thum, R. A. (2016). Potential utility of environmental DNA for early detection of Eurasian watermilfoil. Journal of Aquatic Plant Management, 54, 46–49.

Niemeyer, B., Epp, L. S., Stoof-Leichsenring, K. R., Pestryakova, L. A., & Herzschuh, U. (2017). A comparison of sedimentary DNA and pollen from lake sediments in recording vegetation composition at the Siberian treeline. Molecular Ecology Resources, 17(6), e46–e62. https://doi.org/10.1111/1755-0998.12689

O’Hare, M. T., Baattrup-Pedersen, A., Baumgarte, I., Freeman, A., Gunn, I. D. M., Lázár, A. N., Sinclair, R., Wade, A. J., & Bowes, M. J. (2018). Responses of Aquatic Plants to Eutrophication in Rivers: A Revised Conceptual Model. Frontiers in Plant Science, 9. https://www.frontiersin.org/article/10.3389/fpls.2018.00451

Ortega, A., Geraldi, N. R., Díaz-Rúa, R., Ørberg, S. B., Wesselmann, M., Krause-Jensen, D., & Duarte, C. M. (2020). A DNA mini-barcode for marine macrophytes. Molecular Ecology Resources, 20(4), 920–935. https://doi.org/10.1111/1755-0998.13164

Ottoni, C., Borić, D., Cheronet, O., Sparacello, V., Dori, I., Coppa, A., Antonović, D., Vujević, D., Price, T. D., Pinhasi, R., & Cristiani, E. (2021). Tracking the transition to agriculture in Southern Europe through ancient DNA analysis of dental calculus. Proceedings of the National Academy of Sciences, 118(32), e2102116118. https://doi.org/10.1073/pnas.2102116118

Palacios Mejia, M., Curd, E., Edalati, K., Renshaw, M. A., Dunn, R., Potter, D., Fraga, N., Moore, J., Saiz, J., Wayne, R., & Parker, S. S. (2021). The utility of environmental DNA from sediment and water samples for recovery of observed plant and animal species from four Mojave Desert springs. Environmental DNA, 3(1), 214–230. https://doi.org/10.1002/edn3.161

Pansu, J., Winkworth, R. C., Hennion, F., Gielly, L., Taberlet, P., & Choler, P. (2015). Long-lasting modification of soil fungal diversity associated with the introduction of rabbits to a remote sub-Antarctic archipelago. Biology Letters, 11(9), 20150408. https://doi.org/10.1098/rsbl.2015.0408

Parducci, L., Alsos, I. G., Unneberg, P., Pedersen, M. W., Han, L., Lammers, Y., Salonen, J. S., Väliranta, M. M., Slotte, T., & Wohlfarth, B. (2019). Shotgun Environmental DNA, Pollen, and Macrofossil Analysis of Lateglacial Lake Sediments From Southern Sweden. Frontiers in Ecology and Evolution, 7, 189. https://doi.org/10.3389/fevo.2019.00189

Parducci, L., Bennett, K. D., Ficetola, G. F., Alsos, I. G., Suyama, Y., Wood, J. R., & Pedersen, M. W. (2017). Ancient plant DNA in lake sediments. New Phytologist, 214(3), 924–942. https://doi.org/10.1111/nph.14470

Parducci, L., Matetovici, I., Fontana, S. L., Bennett, K. D., Suyama, Y., Haile, J., Kjær, K. H., Larsen, N. K., Drouzas, A. D., & Willerslev, E. (2013). Molecular- and pollen-based vegetation analysis in lake sediments from central Scandinavia. Molecular Ecology, 22(13), 3511–3524. https://doi.org/10.1111/mec.12298

Parducci, L., Väliranta, M., Salonen, J. S., Ronkainen, T., Matetovici, I., Fontana, S. L., Eskola, T., Sarala, P., & Suyama, Y. (2015). Proxy comparison in ancient peat sediments: Pollen, macrofossil and plant DNA. Philosophical Transactions of the Royal Society B: Biological Sciences, 370(1660), 20130382. https://doi.org/10.1098/rstb.2013.0382

Peabody, M. A., Van Rossum, T., Lo, R., & Brinkman, F. S. L. (2015). Evaluation of shotgun metagenomics sequence classification methods using in silico and in vitro simulated communities. BMC Bioinformatics, 16(1), 362. https://doi.org/10.1186/s12859-015-0788-5

Pedersen, M. W., Ginolhac, A., Orlando, L., Olsen, J., Andersen, K., Holm, J., Funder, S., Willerslev, E., & Kjær, K. H. (2013). A comparative study of ancient environmental DNA to pollen and macrofossils from lake sediments reveals taxonomic overlap and additional plant taxa. Quaternary Science Reviews, 75, 161–168. https://doi.org/10.1016/j.quascirev.2013.06.006

Pedersen, M. W., Ruter, A., Schweger, C., Friebe, H., Staff, R. A., Kjeldsen, K. K., Mendoza, M. L. Z., Beaudoin, A. B., Zutter, C., Larsen, N. K., Potter, B. A., Nielsen, R., Rainville, R. A., Orlando, L., Meltzer, D. J., Kjær, K. H., & Willerslev, E. (2016). Postglacial viability and colonization in North America’s ice-free corridor. Nature, 537(7618), 45–49. https://doi.org/10.1038/nature19085

Penning, W. E., Mjelde, M., Dudley, B., Hellsten, S., Hanganu, J., Kolada, A., van den Berg, M., Poikane, S., Phillips, G., Willby, N., & Ecke, F. (2008). Classifying aquatic macrophytes as indicators of eutrophication in European lakes. Aquatic Ecology, 42(2), 237–251. https://doi.org/10.1007/s10452-008-9182-y

Poikane, S., Portielje, R., Denys, L., Elferts, D., Kelly, M., Kolada, A., Mäemets, H., Phillips, G., Søndergaard, M., Willby, N., & van den Berg, M. S. (2018). Macrophyte assessment in European lakes: Diverse approaches but convergent views of ‘good’ ecological status. Ecological Indicators, 94, 185–197. https://doi.org/10.1016/j.ecolind.2018.06.056

Powell, C., Malpas, J., Tollett, M., Anderson, D., Dorrington, E., Frye, P., Sieck-Hill, F., & Kunz, K. (2022). *CalFlora*. CalFlora. https://www.calflora.org/

R Core Team. (2022). R: A Language and Environment for Statistical Computing (4.1.1). R Foundation for Statistical Computing. https://www.R-project.org/

Reitsema, R. E., Meire, P., & Schoelynck, J. (2018). The Future of Freshwater Macrophytes in a Changing World: Dissolved Organic Carbon Quantity and Quality and Its Interactions With Macrophytes. Frontiers in Plant Science, 9. https://www.frontiersin.org/article/10.3389/fpls.2018.00629

Rijal, D. P., Heintzman, P. D., Lammers, Y., Yoccoz, N. G., Lorberau, K. E., Pitelkova, I., Goslar, T., Murguzur, F. J. A., Salonen, J. S., Helmens, K. F., Bakke, J., Edwards, M. E., Alm, T., Bråthen, K. A., Brown, A. G., & Alsos, I. G. (2021). Sedimentary ancient DNA shows terrestrial plant richness continuously increased over the Holocene in northern Fennoscandia. In Science Advances (Vol. 7, Issue eabf9557, p. 16).

Sawada, M., Viau, A. E., & Gajewski, K. (2003). The biogeography of aquatic macrophytes in North America since the Last Glacial Maximum. Journal of Biogeography, 30(7), 999–1017. https://doi.org/10.1046/j.1365-2699.2003.00866.x

Sawafuji, R., Saso, A., Suda, W., Hattori, M., & Ueda, S. (2020). Ancient DNA analysis of food remains in human dental calculus from the Edo period, Japan. PLOS ONE, 15(3), e0226654. https://doi.org/10.1371/journal.pone.0226654

Schabacker, J. C., Amish, S. J., Ellis, B. K., Gardner, B., Miller, D. L., Rutledge, E. A., Sepulveda, A. J., & Luikart, G. (2020). Increased eDNA detection sensitivity using a novel high-volume water sampling method. Environmental DNA, 2(2), 244–251. https://doi.org/10.1002/edn3.63

Scriver, M., Marinich, A., Wilson, C., & Freeland, J. (2015). Development of species-specific environmental DNA (eDNA) markers for invasive aquatic plants. Aquatic Botany, 122, 27–31. https://doi.org/10.1016/j.aquabot.2015.01.003

Seersholm, F. V., Pedersen, M. W., Søe, M. J., Shokry, H., Mak, S. S. T., Ruter, A., Raghavan, M., Fitzhugh, W., Kjær, K. H., Willerslev, E., Meldgaard, M., Kapel, C. M. O., & Hansen, A. J. (2016). DNA evidence of bowhead whale exploitation by Greenlandic Paleo-Inuit 4,000 years ago. Nature Communications, 7(1), 13389. https://doi.org/10.1038/ncomms13389

Shackleton, M. E., Rees, G. N., Watson, G., Campbell, C., & Nielsen, D. (2019). Environmental DNA reveals landscape mosaic of wetland plant communities. Global Ecology and Conservation, 19, e00689. https://doi.org/10.1016/j.gecco.2019.e00689

Sheldon, R. B., & Boylen, C. W. (1977). Maximum Depth Inhabited by Aquatic Vascular Plants. The American Midland Naturalist, 97(1), 248–254. https://doi.org/10.2307/2424706

Sjögren, P., Edwards, M. E., Gielly, L., Langdon, C. T., Croudace, I. W., Merkel, M. K. F., Fonville, T., & Alsos, I. G. (2017). Lake sedimentary DNA accurately records 20th Century introductions of exotic conifers in Scotland. New Phytologist, 213(2), 929–941. https://doi.org/10.1111/nph.14199

South African National Biodiversity Institute. (2022). *PlantZAfrica*. PlantZAfrica. http://pza.sanbi.org/

Stoof-Leichsenring, K. R., Huang, S., Liu, S., Jia, W., Li, K., Liu, X., Pestryakova, L. A., & Herzschuh, U. (2022). Sedimentary DNA identifies modern and past macrophyte diversity and its environmental drivers in high-latitude and high-elevation lakes in Siberia and China. Limnology and Oceanography, 9999(n/a), 16. https://doi.org/10.1002/lno.12061

Sønstebø, J. H., Gielly, L., Brysting, A. K., Elven, R., Edwards, M., Haile, J., Willerslev, E., Coissac, E., Rioux, D., Sannier, J., Taberlet, P., & Brochmann, C. (2010). Using next-generation sequencing for molecular reconstruction of past Arctic vegetation and climate. Molecular Ecology Resources, 10(6), 1009–1018. https://doi.org/10.1111/j.1755-0998.2010.02855.x

Tabares, X., Zimmermann, H., Dietze, E., Ratzmann, G., Belz, L., Vieth-Hillebrand, A., Dupont, L., Wilkes, H., Mapani, B., & Herzschuh, U. (2020). Vegetation state changes in the course of shrub encroachment in an African savanna since about 1850 CE and their potential drivers. Ecology and Evolution, 10(2), 962–979. https://doi.org/10.1002/ece3.5955

Taberlet, P., Coissac, E., Pompanon, F., Gielly, L., Miquel, C., Valentini, A., Vermat, T., Corthier, G., Brochmann, C., & Willerslev, E. (2007). Power and limitations of the chloroplast trn L (UAA) intron for plant DNA barcoding. Nucleic Acids Research, 35(3), e14. https://doi.org/10.1093/nar/gkl938

Tela Botanica. (2022). *EFlore*. EFlore. https://www.tela-botanica.org/flore/

ter Schure, A. T. M., Bajard, M., Loftsgarden, K., Høeg, H. I., Ballo, E., Bakke, J., Støren, E. W. N., Iversen, F., Kool, A., Brysting, A. K., Krüger, K., & Boessenkool, S. (2021). Anthropogenic and environmental drivers of vegetation change in southeastern Norway during the Holocene. Quaternary Science Reviews, 270, 107175. https://doi.org/10.1016/j.quascirev.2021.107175

Tringe, S. G., & Rubin, E. M. (2005). Metagenomics: DNA sequencing of environmental samples. Nature Reviews Genetics, 6(11), Article 11. https://doi.org/10.1038/nrg1709

Tsukamoto, Y., Yonezawa, S., Katayama, N., & Isagi, Y. (2021). Detection of Endangered Aquatic Plants in Rapid Streams Using Environmental DNA. Frontiers in Ecology and Evolution, 8. https://www.frontiersin.org/article/10.3389/fevo.2020.622291

Tyler, T., Herbertsson, L., Olofsson, J., & Olsson, P. A. (2021). Ecological indicator and traits values for Swedish vascular plants. Ecological Indicators, 120, 106923. https://doi.org/10.1016/j.ecolind.2020.106923

Tyler, T., & Olsson, P. A. (2016). Substrate pH ranges of south Swedish bryophytes—Identifying critical pH values and richness patterns. Flora, 223, 74–82. https://doi.org/10.1016/j.flora.2016.05.006

Tyrrell, C. D., Chambers, P. A., & Culp, J. M. (2022). Harnessing aquatic plant growth forms to apply European nutrient-enrichment bioindicators to Canadian waters. *Applications in Plant Sciences*, *n/a*(n/a), e11487. https://doi.org/10.1002/aps3.11487

United States Department of Agriculture – National Wetland Plant List. (2022). USDA Plant Database—Wetland Status. https://plants.usda.gov/home/wetlandSearch

Väliranta, M., Kultti, S., Nyman, M., & Sarmaja-Korjonen, K. (2005). Holocene development of aquatic vegetation in shallow Lake Njargajavri, Finnish Lapland, with evidence of water-level fluctuations and drying. Journal of Paleolimnology, 34(2), 203–215. https://doi.org/10.1007/s10933-005-1840-7

Väliranta, M., Salonen, J. S., Heikkilä, M., Amon, L., Helmens, K., Klimaschewski, A., Kuhry, P., Kultti, S., Poska, A., Shala, S., Veski, S., & Birks, H. H. (2015). Plant macrofossil evidence for an early onset of the Holocene summer thermal maximum in northernmost Europe. Nature Communications, 6(1), 6809. https://doi.org/10.1038/ncomms7809

von Hippel, B., Stoof-Leichsenring, K. R., Schulte, L., Seeber, P., Epp, L. S., Biskaborn, B. K., Diekmann, B., Melles, M., Pestryakova, L., & Herzschuh, U. (2022). Long-term fungus–plant covariation from multi-site sedimentary ancient DNA metabarcoding. Quaternary Science Reviews, 295, 107758. https://doi.org/10.1016/j.quascirev.2022.107758

Voss, E. G., & Reznicek, A. A. (2012). Field Manual of Michigan Flora. University of Michigan Press.

Walsh, N. G., & Entwisle, T. J. (1994). Flora of Victoria (Vols. 2–4). Inkata Press.

Wang, Y., Pedersen, M. W., Alsos, I. G., De Sanctis, B., Racimo, F., Prohaska, A., Coissac, E., Owens, H. L., Merkel, M. K. F., Fernandez-Guerra, A., Rouillard, A., Lammers, Y., Alberti, A., Denoeud, F., Money, D., Ruter, A. H., McColl, H., Larsen, N. K., Cherezova, A. A., … Willerslev, E. (2021). Late Quaternary dynamics of Arctic biota from ancient environmental genomics. Nature, 600(7887), 86–92. https://doi.org/10.1038/s41586-021-04016-x

Wang, L., Yang, T., Hei, P., Zhang, J., Yang, J., Luo, T., Zhou, G., Liu, C., Wang, R., & Chen, F. (2022). Internal phosphorus cycling in macrophyte-dominated eutrophic lakes and its implications. Journal of Environmental Management, 306, 114424. https://doi.org/10.1016/j.jenvman.2021.114424

Weigand, H., Beermann, A. J., Čiampor, F., Costa, F. O., Csabai, Z., Duarte, S., Geiger, M. F., Grabowski, M., Rimet, F., Rulik, B., Strand, M., Szucsich, N., Weigand, A. M., Willassen, E., Wyler, S. A., Bouchez, A., Borja, A., Čiamporová-Zaťovičová, Z., Ferreira, S., … Ekrem, T. (2019). DNA barcode reference libraries for the monitoring of aquatic biota in Europe: Gap-analysis and recommendations for future work. Science of The Total Environment, 678, 499–524. https://doi.org/10.1016/j.scitotenv.2019.04.247

WFO: World Flora Online. (2022). *World Flora Online*. World Flora Online. http://www.worldfloraonline.org

Willerslev, E., Davison, J., Moora, M., Zobel, M., Coissac, E., Edwards, M. E., Lorenzen, E. D., Vestergård, M., Gussarova, G., Haile, J., Craine, J., Gielly, L., Boessenkool, S., Epp, L. S., Pearman, P. B., Cheddadi, R., Murray, D., Bråthen, K. A., Yoccoz, N. G., … Taberlet, P. (2014). Fifty thousand years of Arctic vegetation and megafaunal diet. Nature, 506, 47–51.

Willerslev, E., Hansen, A. J., Binladen, J., Brand, T. B., Gilbert, M. T. P., Shapiro, B., Bunce, M., Wiuf, C., Gilichinsky, D. A., & Cooper, A. (2003). Diverse Plant and Animal Genetic Records from Holocene and Pleistocene Sediments. Science, 300, 791–795.

Wisconsin State Herbarium. (2022). *Online Virtual Flora of Wisconsin*. Online Virtual Flora of Wisconsin. https://wisflora.herbarium.wisc.edu/index.php

Wood, J. R., Crown, A., Cole, T. L., & Wilmshurst, J. M. (2016). Microscopic and ancient DNA profiling of Polynesian dog (kurī) coprolites from northern New Zealand. Journal of Archaeological Science: Reports, 6, 496–505. https://doi.org/10.1016/j.jasrep.2016.03.020

Zhang, J., Hei, P., Shang, Y., Yang, J., Wang, L., Yang, T., Zhou, G., & Chen, F. (2021). Internal Nitrogen Cycle in Macrophyte-Dominated Eutrophic Lakes: Mechanisms and Implications for Ecological Restoration. ACS ES&T Water, 1(11), 2359–2369. https://doi.org/10.1021/acsestwater.1c00203

Zimmermann, H. H., Raschke, E., Epp, L. S., Stoof-Leichsenring, K. R., Schwamborn, G., Schirrmeister, L., Overduin, P. P., & Herzschuh, U. (2017). Sedimentary ancient DNA and pollen reveal the composition of plant organic matter in Late Quaternary permafrost sediments of the Buor Khaya Peninsula (north-eastern Siberia). Biogeosciences, 14(3), 575–596. https://doi.org/10.5194/bg-14-575-2017

Zou, Y.-F., Chen, K.-Y., & Lin, H.-J. (2021). Significance of belowground production to the long-term carbon sequestration of intertidal seagrass beds. Science of The Total Environment, 800, 149579. https://doi.org/10.1016/j.scitotenv.2021.149579

