## Supplementary Figure S1 for "Environmental DNA of aquatic macrophytes: the potential for reconstructing past and present vegetation and environments"

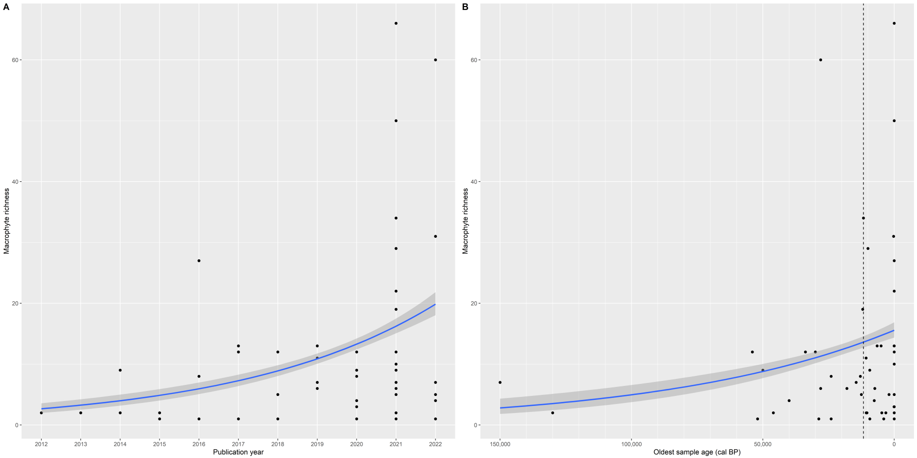


Figure S1: Plots of the Generalised Linear Models investigating how the (A) year of publication and (B) age of the oldest sample affect the total richness of aquatic macrophytes detected. The dashed line indicates the Pleistocene–Holocene border, at 11,700 cal BP.
