## Supplementary figures and images for "Environmental DNA of aquatic macrophytes: the potential for reconstructing past and present vegetation and environments"

### Supplementary Figure S2

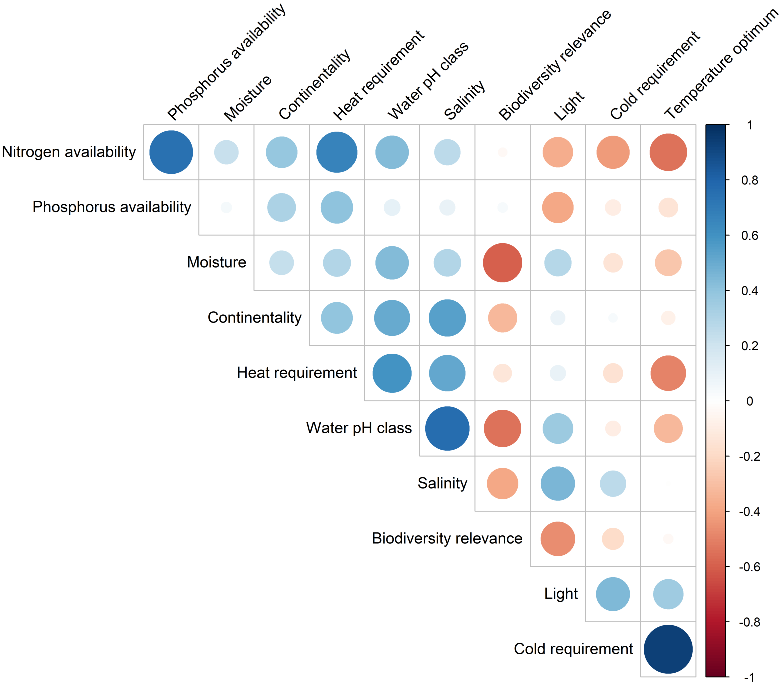


Figure S2: Correlogram of the preselected environmental traits.

### Supplementary Figure S3

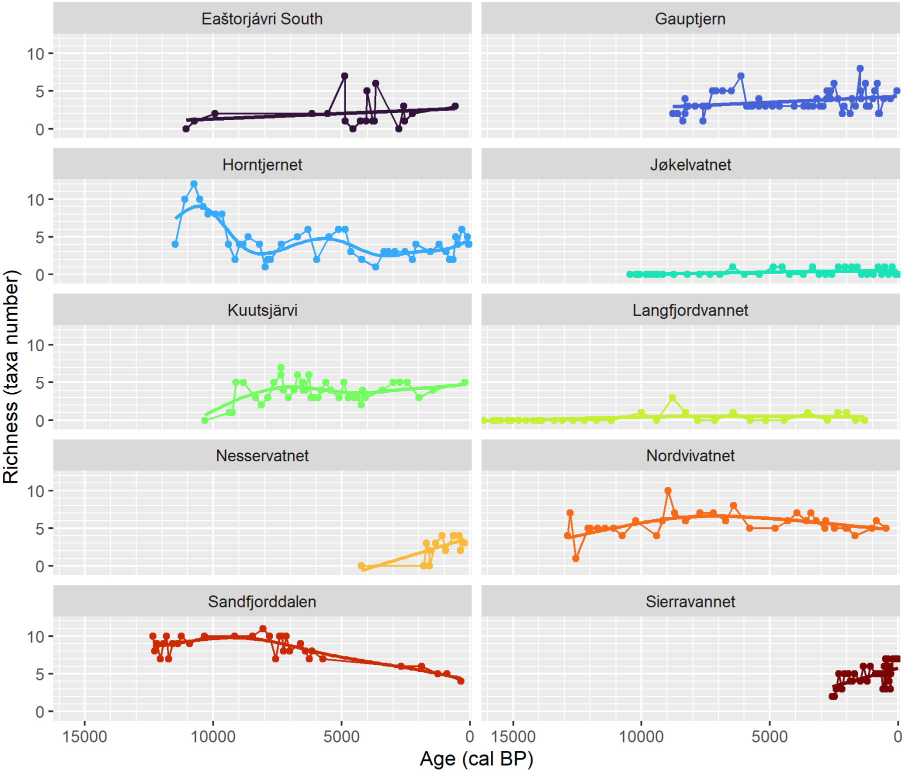


Figure S3: Lake-specific richness in aquatic macrophyte taxa through time.
