## Supplementary Figure S4 for "Environmental DNA of aquatic macrophytes: the potential for reconstructing past and present vegetation and environments"

| | Heat requirement | Cold requirement | Continentality | pH | Nitrogen availability | Light optimum |
|---|---|---|---|---|---|---|
| Callitriche cf. palustris | 3 | 7 | 6 | 6 | 5 | 6 |
| Callitriche hermaphroditica | 5 | 7 | 7 | 7 | 5 | 5 |
| Carex rotundata/rostrata | 3 | (NA) | 5.5 | 3.5 | 3 | (NA) |
| Ceratophyllum demersum | 6 | 1 | 6 | 8 | 9 | 6 |
| Cicuta virosa | 7 | 5 | 7 | 5 | 7 | 5 |
| Comarum palustre | 3 | 4 | 5 | 4 | 4 | 5 |
| Hippuris spp. | 5.67 | (NA) | 6.33 | 6.33 | 5 | 6.67 |
| Isoetes spp. | 3.5 | 8 | 5 | 3.5 | 2 | 7 |
| Menyanthes trifoliata | 3 | 3 | 5 | 5 | 4 | 5 |
| Myriophyllum alterniflorum | 4 | 2 | 5 | 4 | 4 | 5 |
| Myriophyllum sibiricum | 4 | 7 | 7 | 7 | 5 | 7 |
| Nuphar spp. / Nymphaea alba | 5.33 | (NA) | 5.33 | 5.33 | 5.67 | 6 |
| Phragmites australis | 6 | 1 | 6 | 6 | 7 | 5 |
| Potamogeton alpinus | 3 | 5 | 5 | 4 | 6 | 5 |
| Potamogeton berchtoldii/friesii/natans/obtusifolius | 5.75 | 3.5 | 5 | 6.25 | 6 | 6.25 |
| Potamogeton berchtoldii/friesii/obtusifolius | 6 | 4 | 5 | 6.67 | 6.33 | 6 |
| Potamogeton perfoliatus | 4 | 2 | 6 | 7 | 5 | 7 |
| Potamogeton perfoliatus/alpinus/gramineus | 3.67 | 3.67 | 6 | (NA) | 5 | 6 |
| Potamogeton praelongus | 4 | 6 | 6 | 7 | 4 | 7 |
| Ranunculus cf. reptans | 3 | 8 | 5 | 4 | 4 | 6 |
| Ranunculus cf. reptans/flammula | (NA) | (NA) | 4.5 | 4 | 5 | 5.5 |
| Ranunculus peltatus | 4 | 2 | 7 | 6 | 7 | 5 |
| Sparganium angustifolium/hyperboreum | 3.5 | (NA) | 5 | 3.5 | 4 | 6.5 |
| Sparganium erectum/natans/glomeratum/emersum | (NA) | (NA) | 5.25 | (NA) | (NA) | 5.5 |
| Stuckenia filiformis | 5 | 6 | 5 | 8 | 4 | 7 |
| Stuckenia vaginata | 7 | 10 | 6 | 7 | (NA) | 7 |
| Subularia aquatica | 4 | 7 | 6 | 5 | 5 | 7 |
| Utricularia spp. | 3.33 | 4 | 5.67 | (NA) | 1 | 6.67 |

Figure S4: Graphical visualisation of environmental niches per taxon, for the selected traits.
