## Supplementary Figure S6 for "Environmental DNA of aquatic macrophytes: the potential for reconstructing past and present vegetation and environments"

Figure S6: Proportion of traits values over time in PCR replicates for lakes (a) Gauptjern, (b) Sandfjorddalen, (c) Horntjernet, (d) Kuutsjärvi, (e) Nordvivatnet, (f) Sierravannet, (g) Nesservatnet, (h) Eaštorjávri South, (i) Jøkelvatnet, and (j) Langfjordvannet

a) Gauptjern

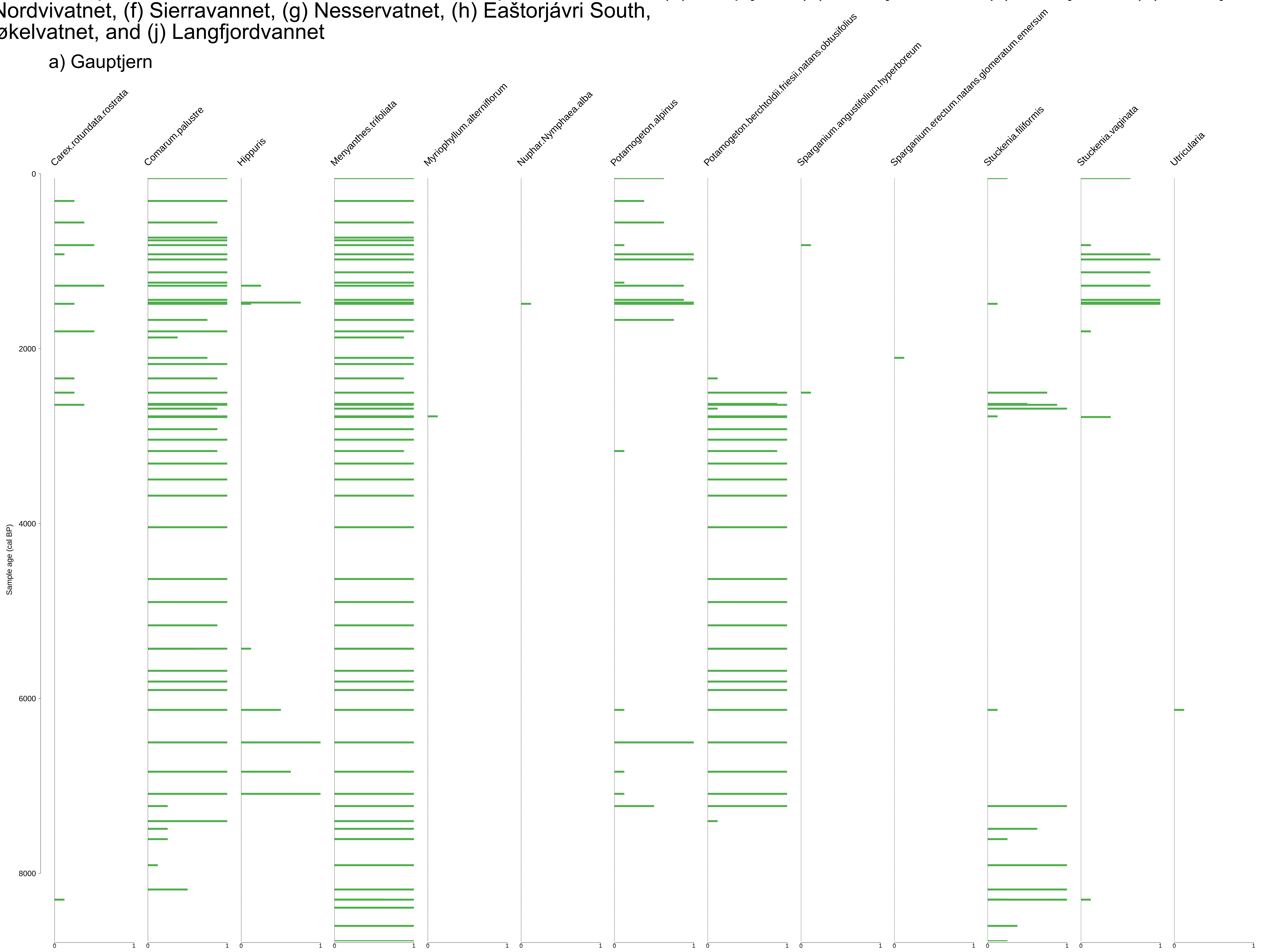

b) Sandfjorddalen

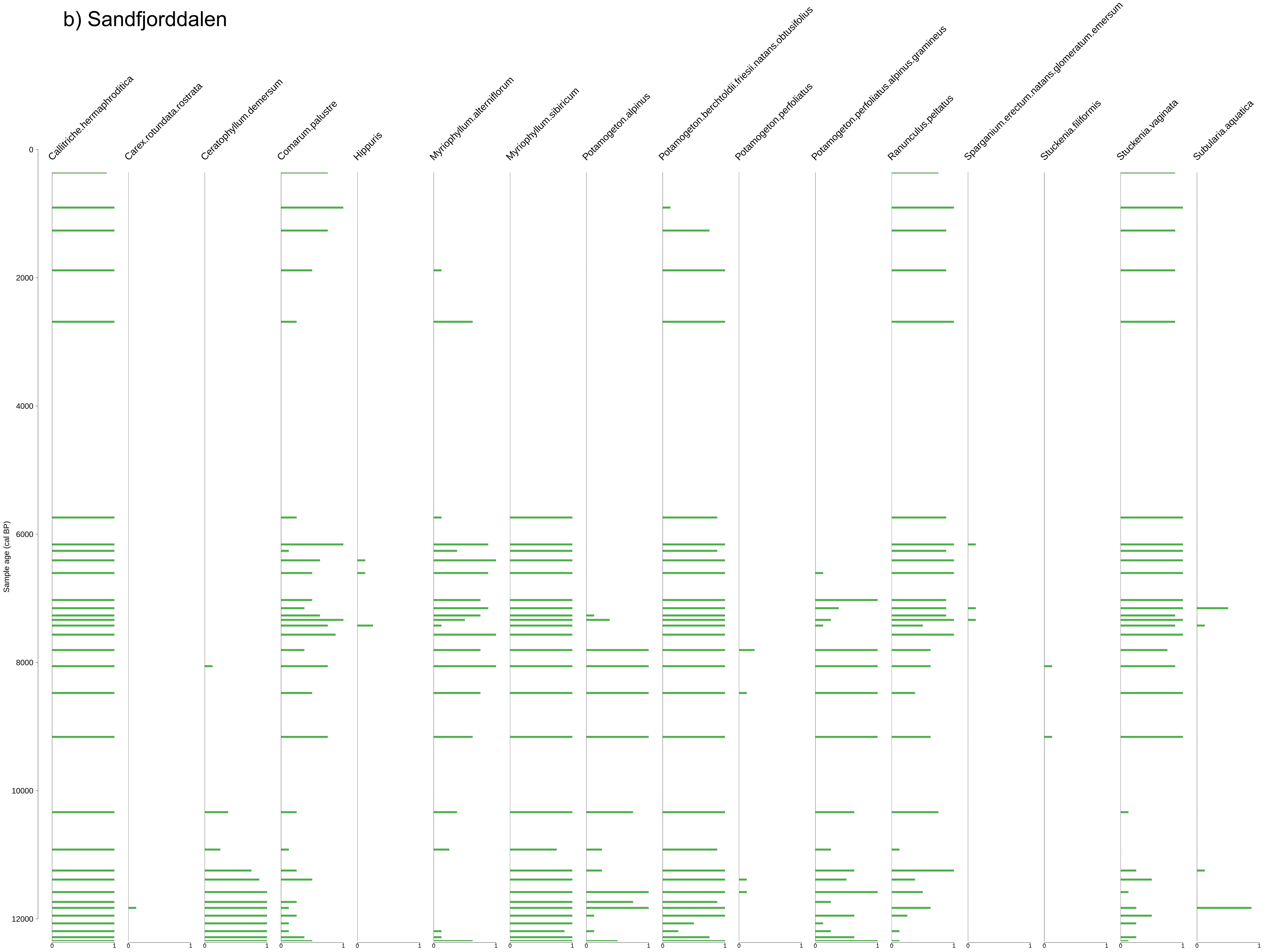

c) Horntjernet

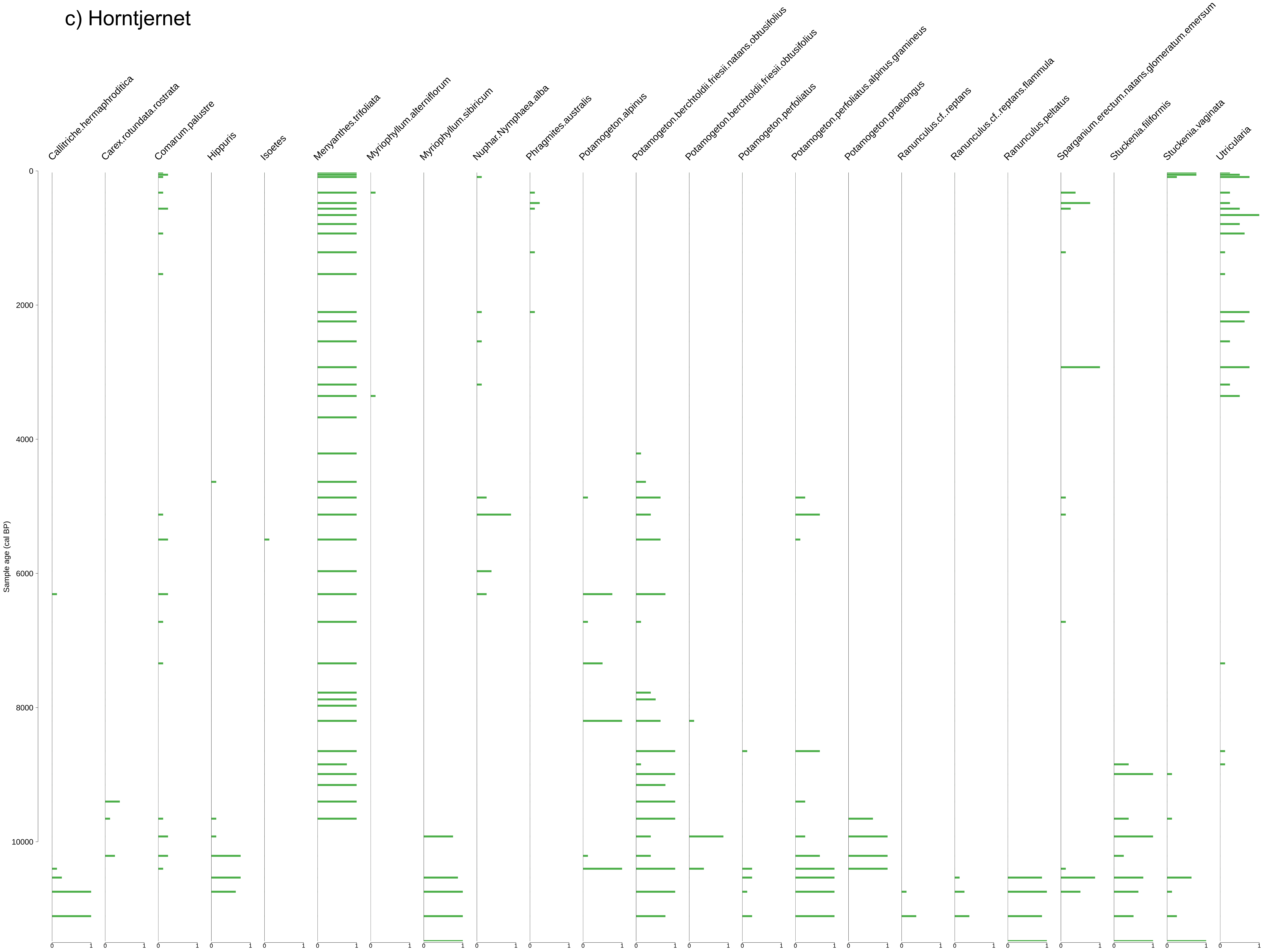

### d) Kuutsjärvi

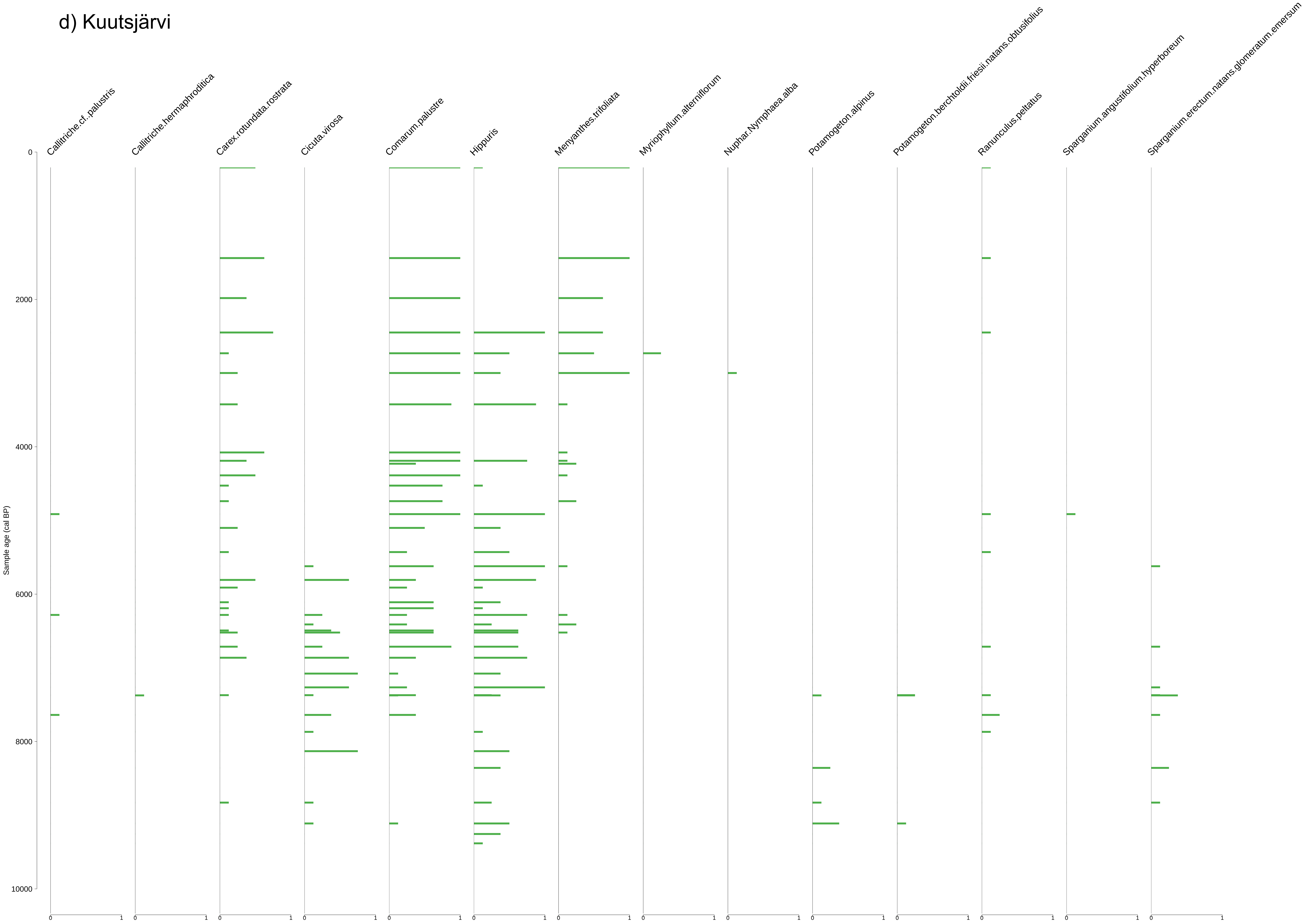

#### e) Eaštorjávri South

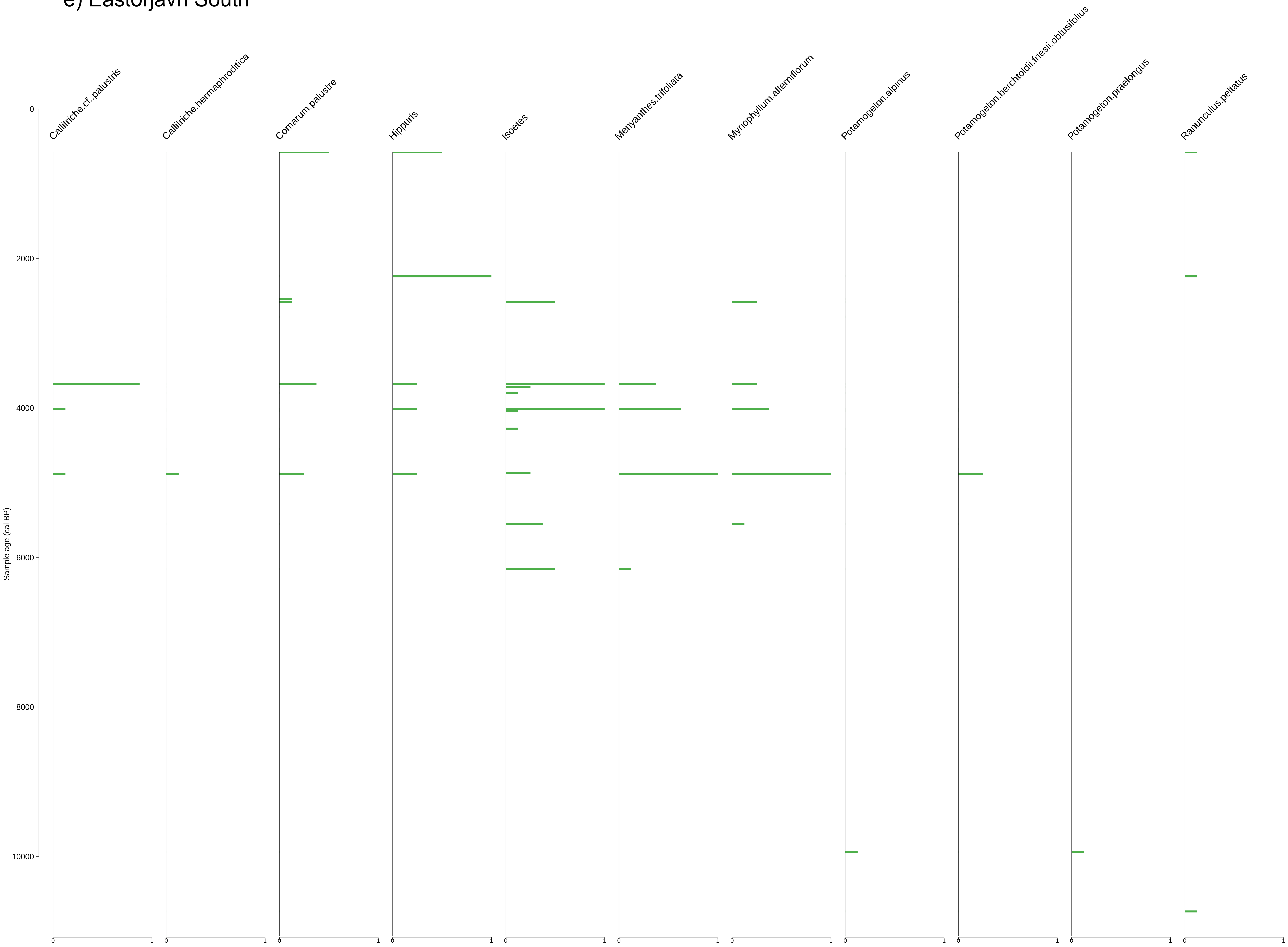

### f) Jøkelvatnet

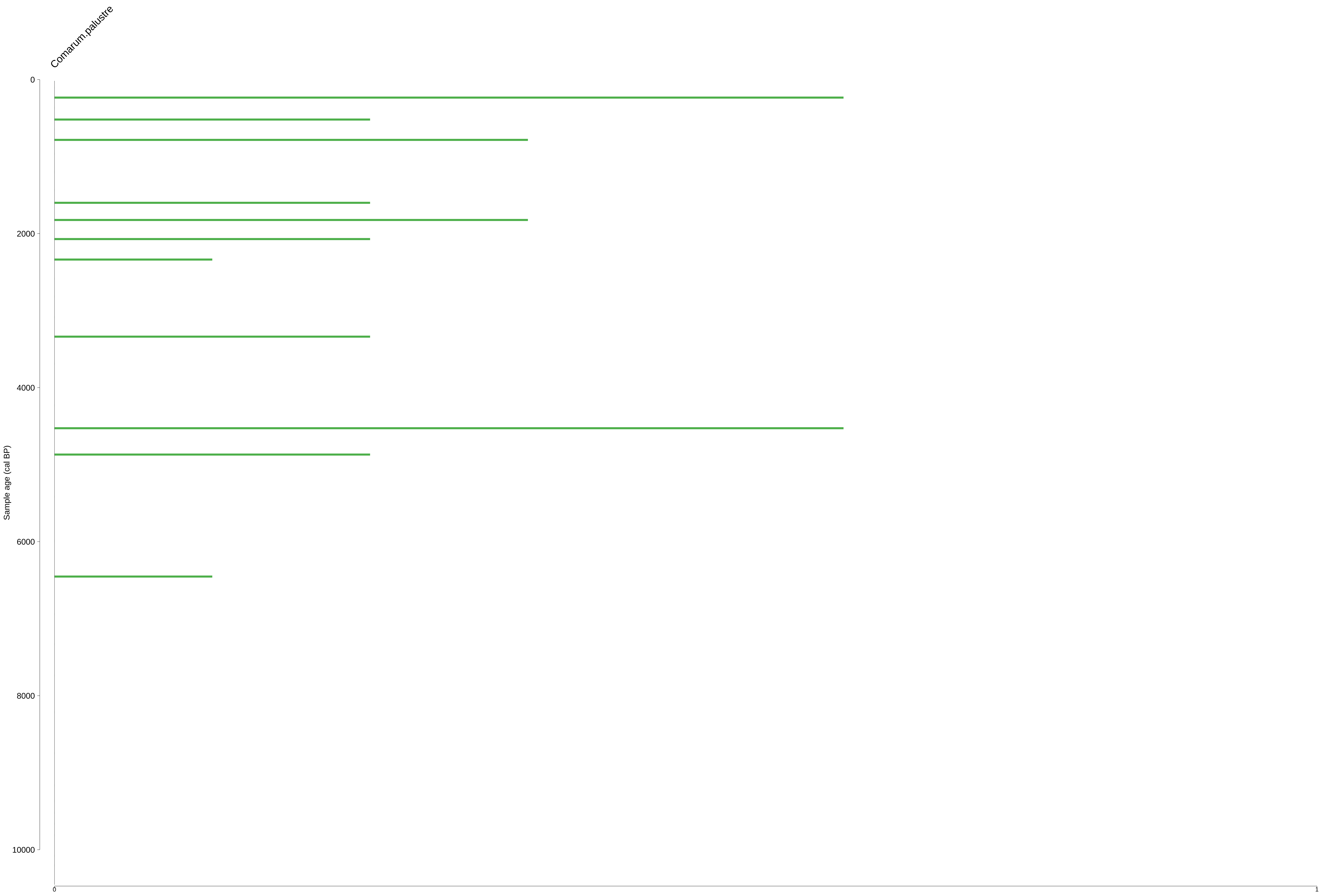

### g) Langfjordvannet

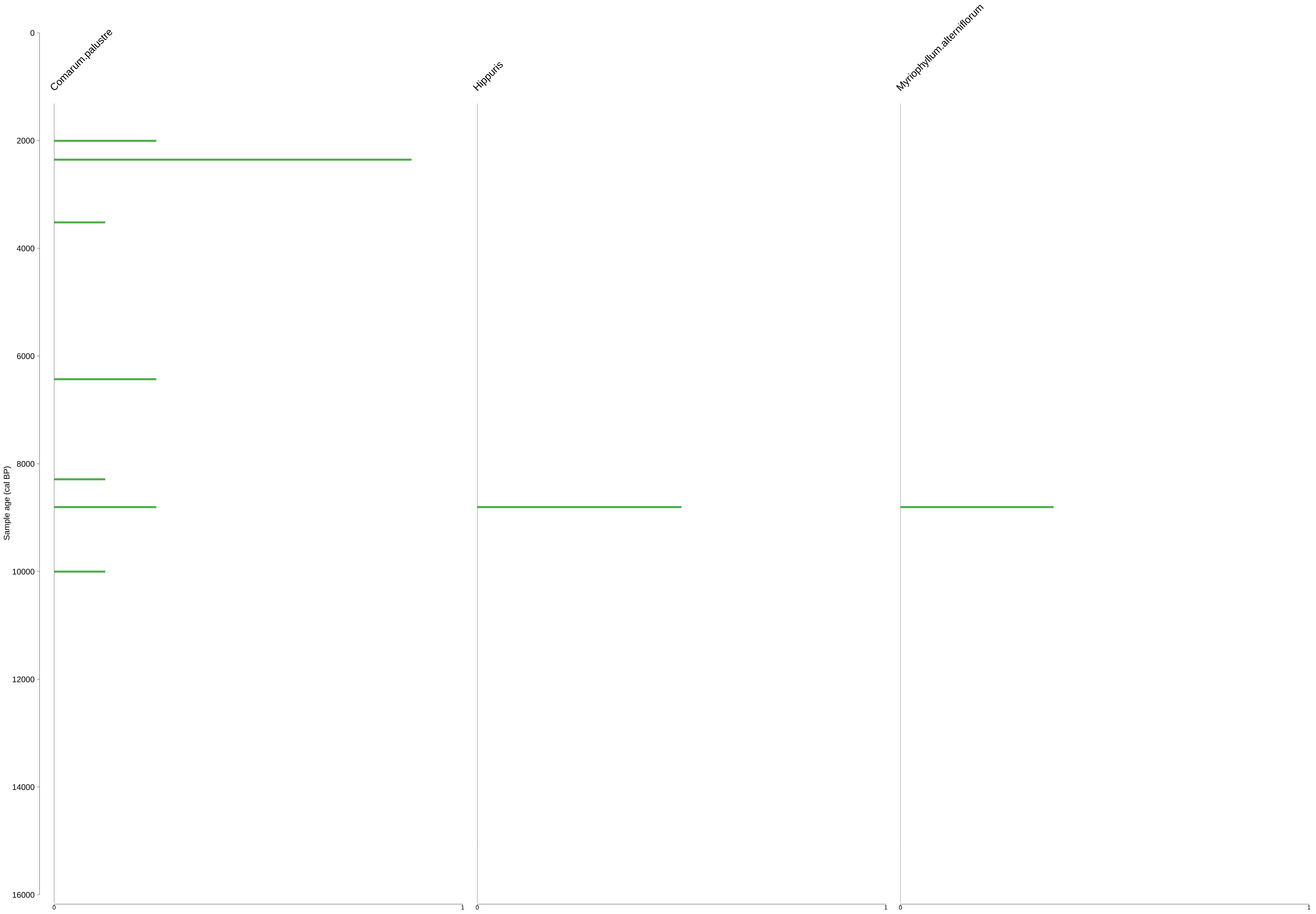

#### h) Nesservatnet

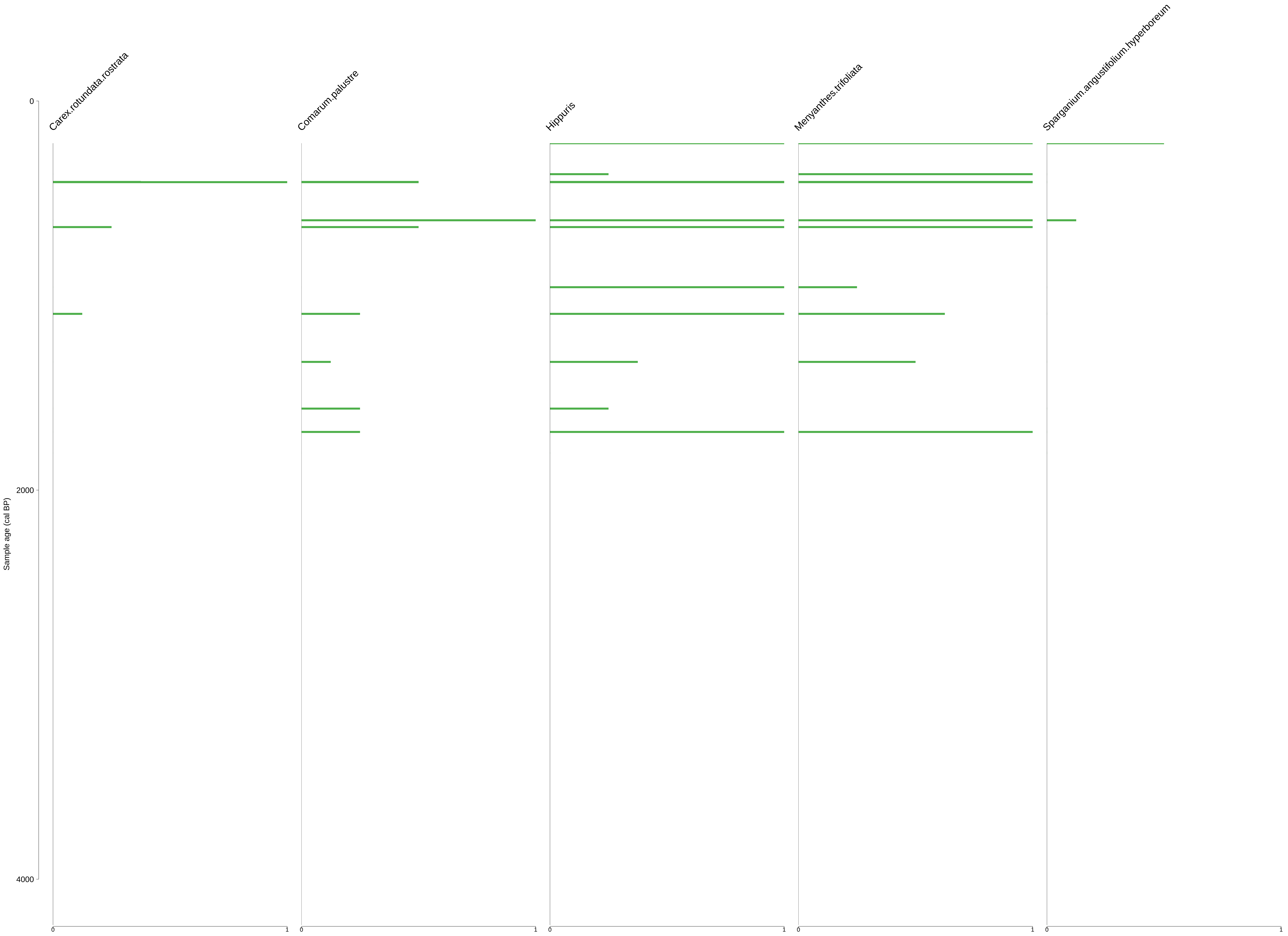

#### i) Nordvivatnet

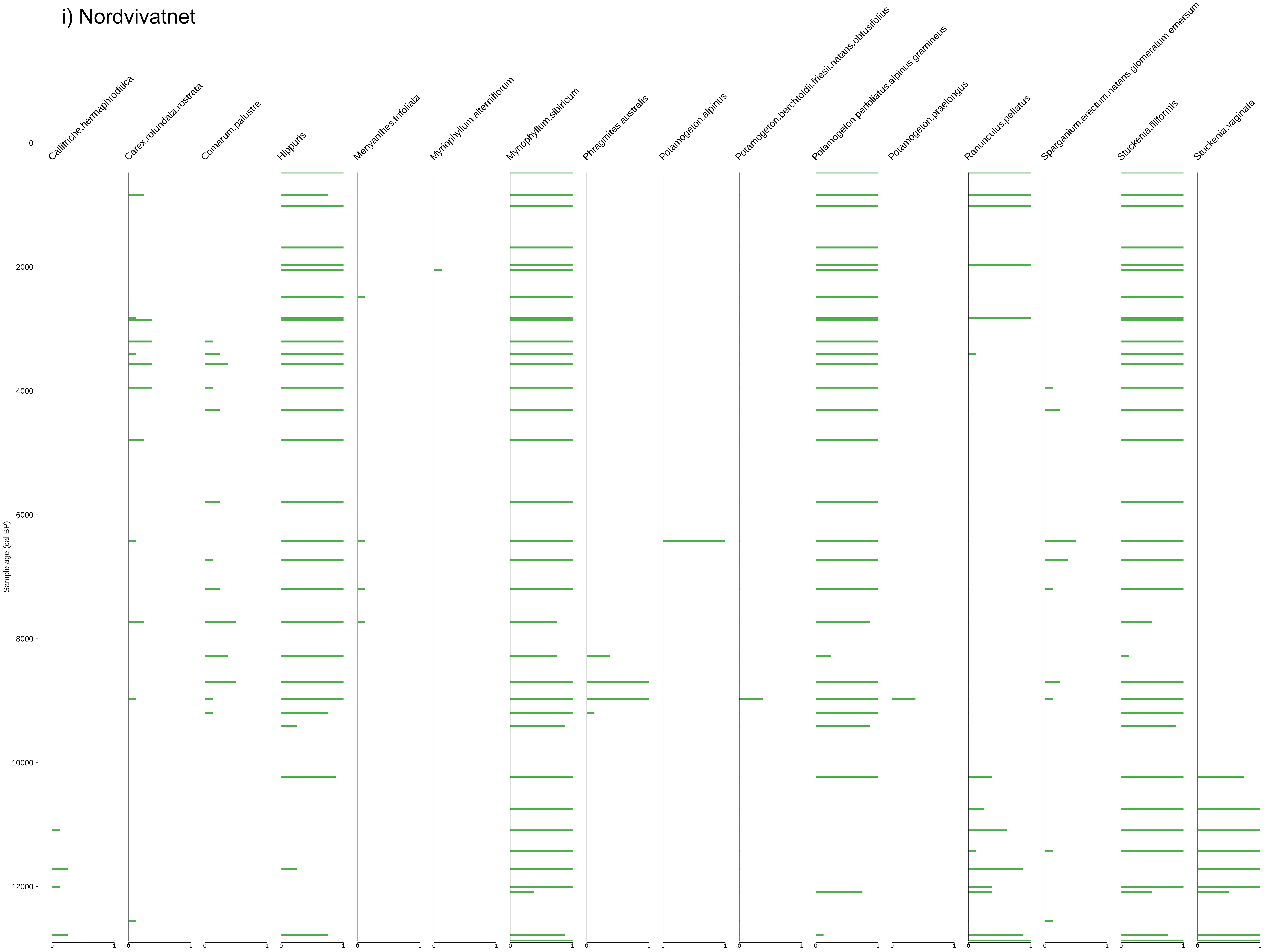

### j) Sierravannet

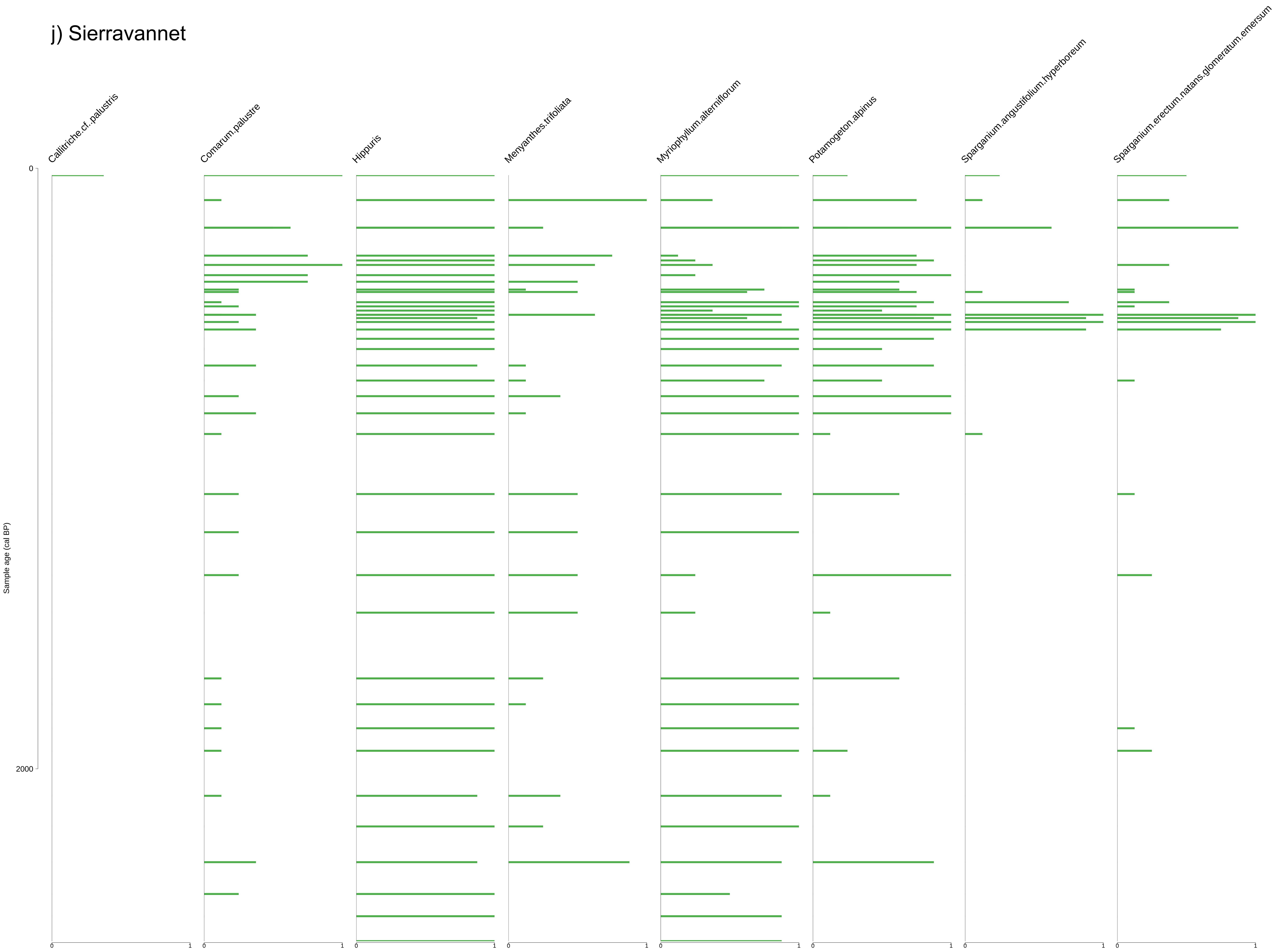
