## Supplementary Table S4 for "Environmental DNA of aquatic macrophytes: the potential for reconstructing past and present vegetation and environments"

Table S4. A comparison table for the Tyler et al. (2021) traits database as derived from Ellenberg et al. (2001) and commonly used aquatic plant life-form classifications. The maximum depth for submerged aquatic plants ^1^ is based on Sheldon and Boylen (1977) but will vary with species and water quality.

| **Tyler Code** | **Tyler Desc.** | **Life-form** | **Raunkiaer life-forms** | **Exemplar Spp.** |
| --- | --- | --- | --- | --- |
| 12 | Deep (> 0.5m) permanent water | Free-floating, Floating-leaved, Submerged (up to 12 m^1^) | Hydrophytes (water plants) | Algae*, Lobelia dortmanna, Nymphaea alba, Potamogeton praelongus, Elodea canadensis* |
| 11 | Shallow (<0.5 m) permanent water | Submerged, Free-floating, Floating-leaved |  | *Callitriche hermaphroditica,*  *Myriophyllum alterniflorum, Elodea canadensis* |
| 10 | Temporarily inundated | Emergent, Marginal | Heliophytes (winter buds under water flowers above water) | *Calla palustris, Carex rostrata, Iris pseudacorus* |
| 9 | Wet-temporarily inundated | Emergent, Marginal |  | *Caltha palustris, Lysimachia thyrsiflora, Ranunculus flammula* |

Ellenberg, H., Weber, H.E., Düll, R., Wirth, V., Werner, W., 2001. Zeigerwerte von Pflanzen in Mitteleuropa (in German with English summary). *Scripta Geobotanica* 18.

Sheldon, R.B. and Boylen, C.W. 1977. Maximum Depth Inhabited by Aquatic Vascular Plants. The American Midland Naturalist 97, 248-254.
