## Supplementary Table S5 for "Environmental DNA of aquatic macrophytes: the potential for reconstructing past and present vegetation and environments"

Taxonomic level

Method	Species	Genus	Family	higher

Targeted capture	0*.*2881	0*.*2881	0*.*0603	0*.*0603

Metabarcoding	* 2*.*261	10*^−^*^6^	* 0*.*0497	* 1*.*169	10*^−^*^7^	0*.*1341

*×	×*

Shotgun metagenomics	1	0*.*5389	0*.*4331	0*.*7104

Table S5: Wilcoxon test *p*-values evaluating the difference in taxonomic reso- lution between aquatic taxa and all vascular plants, for each methodological approach. Asterisks indicate a significant (*p <* 0.05) difference. Values were rounded to four decimal places.
